# Title: Single-molecule diffusion-based estimation of ligand effects on G protein-coupled receptors

**DOI:** 10.1101/205161

**Authors:** Masataka Yanagawa, Michio Hiroshima, Yuichi Togashi, Mitsuhiro Abe, Takahiro Yamashita, Yoshinori Shichida, Masayuki Murata, Masahiro Ueda, Yasushi Sako

## Abstract

G protein-coupled receptors (GPCRs) are major drug targets and have high potential for drug discovery. The development of a method for measuring the activities of GPCRs is essential for pharmacology and drug screening. However, it is difficult to measure the effects of a drug by monitoring the receptor on the cell surface, and changes in the concentrations of downstream signaling molecules, which depend on signaling pathway selectivity of the receptor, are used as an index of the receptor activity. Here, we show that single-molecule imaging analysis provides an alternative method for assessing ligand effects on GPCR. We monitored the dynamics of the diffusion of metabotropic glutamate receptor 3 (mGluR3), a class C GPCR, under various ligand conditions by using total internal reflection fluorescence microscopy (TIRFM). The single-molecule tracking analysis demonstrates that changes in the average diffusion coefficient of mGluR3 quantitatively reflect the ligand-dependent activity. Then, we reveal that the diffusion of receptor molecules is altered by the common physiological events associated with GPCRs, including G protein binding or accumulation in clathrin-coated pits, by inhibition experiments and dual-color single-molecule imaging analysis. We also confirm the generality of agonist-induced diffusion change in class A and B GPCRs, demonstrating that the diffusion coefficient is a good index for estimating the ligand effects on many GPCRs regardless of the phylogenetic groups, chemical properties of the ligands, and G protein-coupling selectivity.

**One Sentence Summary:** Single-molecule imaging for evaluating ligand effects on GPCRs by monitoring the diffusion dynamics on the cell surface.

## Introduction

G protein-coupled receptors (GPCRs) constitute the largest superfamily of human membrane proteins, and are classified into several families based on their sequence similarity (1, 2). About 33% of all small-molecule drugs target just 6% of the ~800 human GPCRs (3, 4); thus, GPCRs have immense potential for drug discovery. However, it is difficult to measure the effects of a drug by monitoring the receptor on the cell surface, and changes in the concentrations of downstream signaling molecules, including second messengers, are monitored as an index of the receptor activity (5). These conventional methods require background knowledge about the signaling pathways, including coupling specificity to G protein subtypes. Here, we developed an alternative method for assessing ligand effects on GPCRs by monitoring the movements of receptor molecules on living cells under a microscope.

Total internal reflection fluorescence microscopy (TIRFM) is a common imaging method for observing single molecules on the bottom plasma membrane of a living cell (6–8). Dimerization and diffusion of the M1 muscarinic receptor (9) and the N-formyl-peptide receptor (10) have been measured by TIRFM, but there was limited information about the activation process because the fluorescent dyes were conjugated with agonists in these studies, preventing observation of the inactive state. Recent studies have reported that the oligomerization and diffusion of the adrenergic receptors (11, 12), GABAB receptors (11), and dopamine D2 receptors (13) change upon ligand stimulation; however, the physiological background and generality of these observations are unknown.

Here, we examined the relationship between the diffusion and functional states of metabotropic glutamate receptor 3 (mGluR3) as a model class C GPCR. Class C GPCRs have a large extracellular ligand-binding domain (ECD) on the N-terminal side of the seven alpha-helical transmembrane domains (TMDs) (Fig. 1A). The ECDs function as an obligatory dimer, where dimeric reorientation occurs upon ligand binding (14, 15). The conformational change in ECDs promotes the dimeric rearrangement of TMDs (16–18), activating a protomer of the TMD dimer (19, 20). Single-molecule tracking (SMT) analysis demonstrated that the average diffusion coefficient (*D_Av_*) of mGluR3 quantitatively reflects receptor activity. The inhibition experiments and dual-color TIRFM analysis indicated that the slowing of mGluR3 was related to the decoupling of the receptor/G protein coupling complex followed by the receptors accumulating in clathrin-coated pits (CCPs). We verified the generality of the agonist-induced change in the diffusion dynamics of GPCRs by comparing the *D_Av_* of nine GPCRs in various phylogenetic positions.

**Fig. 1.**
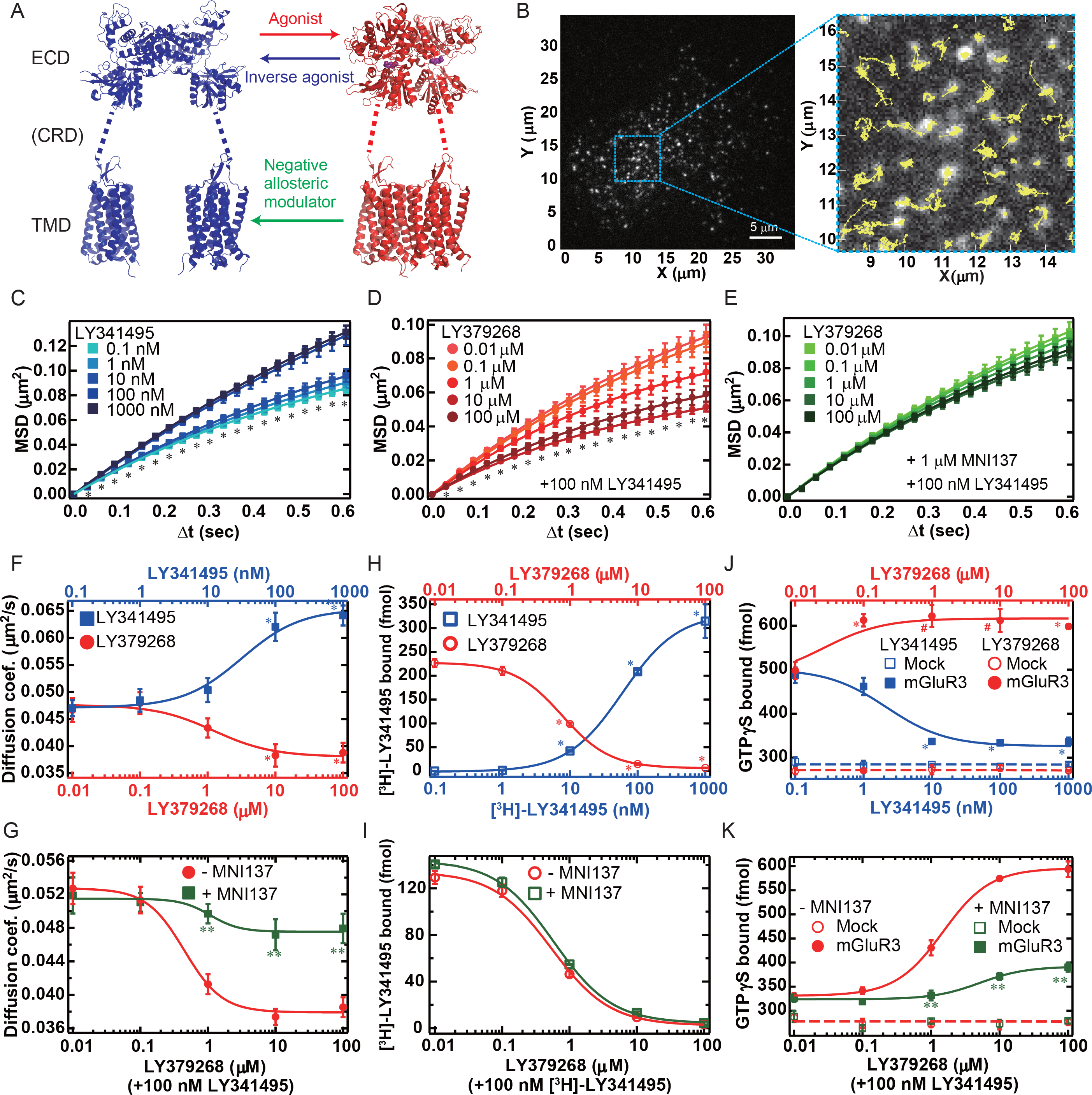
Activation model, TIRFM image, MSD-Δt plots, and comparison of diffusion, ligand occupancy, and G protein activation of mGluR3. **(A)** Activation model of mGluR. The crystal structures of ECDs of mGluR1 in the inactive (blue: 1EWT) and active states (red: 1EWK) are constructed with PyMol (http://www.pymol.org/). The crystal structure of the TMD (blue and red: 4OR2) is also shown. **(B)** Example TIRFM image of HEK293 cell expressing TMR-labeled mGluR3 (left panel: whole image, right panel: enlarged view of the blue dashed square in the left panel). The trajectories of mGluR3 molecules are shown as yellow lines in the right panel. **(C-E)** MSD-Δ*t* plots of the trajectories of mGluR3 under various ligand conditions. Inverse agonist (LY341495) dependency is shown in (C). Agonist (LY379268) dependencies at 100 nM LY314195 without (D) and with (E) 1 μM MNI137. All the data are shown as mean ± SEM (*n* = 20 cells). * Significant difference in MSD among five ligand conditions at each Δ*t* (*p* < 0.01; one-way ANOVA). **(F, G)** Dose-dependent changes of *D_Av_*. LY341495- (blue squares, EC_50_: 28.2 ± 0.9 nM) and LY379268- (red circles, EC_50_: 1.19 ± 0.02 μM) dependency without other ligands are shown in (F). LY379268-dependencies at 100 nM LY341495 without (red circles, EC_50_: 1.03 ± 0.08 μM) and with (green squares, EC_50_: 0.34 ± 0.003 μM) 1 μM MNI137 are shown in (G). All the data are shown as mean ± SEM (*n* = 20 cells). * Significant difference compared with the leftmost point in each curve in (F) (*p* < 0.01; t-test, two-tailed). ** Significant difference with and without 1 μM MNI137 in (G) (*p* < 0.01; t-test, two-tailed). **(H, I)** Dose-dependent changes of specific [^3^H]-LY341495 binding. [^3^H]-LY341495 saturation binding (H, blue squares, K_d_: 47.4 ± 1.7 nM). Replacement of 100 nM [^3^H]-LY341495 with LY379268 in the absence (H and I, red circles, IC_50_: 0.55 ± 0.08 μM) and presence (I, green squares, IC_50_: 0.60 ± 0.03 μM) of 1 μM MNI137. The amount of non-specifically bound 100 nM [^3^H]-LY341495 was 50 ± 11 fmol. All the data are shown as mean ± SEM (*n* = 3 independent experiments). The same mGluR3-transfected cell membrane preparation was analyzed within the same panel. * Significant difference compared with the leftmost point in each curve in (H) (*p* < 0.01; t-test, two-tailed). No significant difference was detected with and without 1 μM MNI137 in (I) (*p* > 0.05; t-test, two-tailed). **(J, K)** Dose-dependent changes in G protein activation efficiency of mGluR3-transfected cell membrane. LY341495- (blue closed squares, IC_50_: 2.11 ± 0.18 nM) and LY379268- (red closed circles, EC_50_: 0.025 ± 0.0029 μM) dependencies without other ligands are shown in (J). LY379268-dependencies at 100 nM LY341495 without (red closed circles, EC_50_: 1.77 ± 0.39 μM) and with (green closed squares, EC_50_: 9.34 ± 4.44M) 1 μM MNI137 are shown in (K). Open circles and squares indicate G protein activation efficiency of mock-transfected cell membrane under the same ligand conditions as for closed circles and squares, respectively. All the data are shown as mean ± SEM (*n* = 3-5 independent experiments). *, # Significant differences compared with the leftmost point in each curve in (J) (*p* < 0.01 and p < 0.03, respectively; t-test, two-tailed). ** Significant difference with and without 1 μM MNI137 in (K) (*p* < 0.01; t-test, two-tailed).

## Results

### Expression, fluorescence labeling, single-molecule imaging of HaloTag fusion mGluR3

To determine the relationship between the diffusion coefficient and activity of mGluR3, we monitored the single-molecule movement of tetramethylrhodamine (TMR)-labeled HaloTag fusion mGluR3 on HEK293 cells under various ligand conditions (Fig. 1, Movie 1). The fusion of HaloTag to the C-terminus of mGluR3 did not alter the dimerization, ligand binding, and G protein activation (Fig. S1).

In previous studies, N-terminally SNAP-tagged GPCRs were labeled with non-membrane-permeable fluorophores (11–13). In this method, only the receptor molecules on the cell surface are labeled at a certain time point, similar to a pulse-chase experiment. Therefore, exocytosis and endocytosis of the receptor molecules after labeling alter the composition of visible receptor molecules on the cell surface depending on the incubation time. Especially, agonist-induced internalization of the receptors causes selective loss of the activated receptors. In contrast, we used membrane-permeable fluorophores, which allow uniform labeling of the receptor molecules in a cell. Whole-cell labeling is important for monitoring the total number of receptor molecules, with and without ligand, on the cell surface, including newly exocytosed receptors after labeling.

When using membrane-permeable fluorophores, non-specific binding of the fluorophores in the cell should be evaluated carefully. We used the HaloTag TMR ligand (TMR), HaloTag STELLA Fluor 650 ligand (SF650), and SNAP-Cell 647-SiR ligand (SiR) to label GPCRs or G proteins. TMR showed the least non-specific binding in mock transfected cells (Fig. S2A-F). It was difficult to use more than 300 nM SF650 or SiR for single-molecule imaging because of the high amount of non-specific binding (Fig. S2D-F). We also evaluated the specific binding affinity of fluorophores to Halo/SNAP-tagged proteins based on the difference in fluorescence intensity between the tagged protein-expressing and non-expressing cells (Fig. S2A-C). The affinity to the HaloTag was fourfold higher for SF650 than for TMR. In terms of photostability and brightness, SF650 also performed better than TMR under our experimental conditions (Fig. S2G and H, Materials and Methods). However, TMR achieved a higher rate of dye labeling under low non-specific binding conditions.

In the single-molecule imaging of mGluR3, ~95% of the HaloTag-fused mGluR3 molecules were specifically labeled with 300 nM TMR, according to the saturation binding assay using the HaloTag TMR ligand (Fig. S2A, Materials and Methods). Because HEK293 cells express no detectable mRNA for mGluRs (21), almost all receptor molecules were labeled in the present measurements. The mean density of receptor molecules on a cell surface was 0.40 ± 0.11 particles/ μm2 in the single-molecule images (Fig. S2I; mean ± SD).

### Time-dependent mean square displacement (*MSD*-*Δt*) analysis of HaloTag fusion mGluR3

MSD-Δt plot analysis is often used to evaluate the diffusion coefficient and diffusion mode of membrane proteins (7, 9, 11, 13). The MSD corresponds to the average squared distance that a receptor molecule travels from its starting point within a certain time interval Δt, which is proportional to the apparent lateral diffusion coefficient. A linear MSD-Δt plot is observed when the receptor molecules exhibit simple Brownian diffusion (22). On the other hand, a concave-up or -down shape of the MSD-Δt plot suggests that the receptor molecules exhibit directed or confined diffusion modes, respectively (22).

We quantified the MSD from the trajectories traced by SMT in each cell (Fig. 1B), and analyzed the dose-dependent change in the total average MSDs of the trajectories (Fig. 1C-E). Stimulation with the inverse agonist LY341495 significantly increased the MSD of mGluR3 molecules (Fig. 1C). In contrast, stimulation with the agonist LY379268 significantly decreased the MSD in a dose-dependent manner (Fig. 1D). We also analyzed the LY379268-dependent diffusion change of mGluR3 in the presence of 1 ΔM MNI137, a negative allosteric modulator (NAM) (23). MNI137 binding to the TMD suppressed the agonist-dependent decrease of MSD, and no significant difference was observed upon LY379268 stimulation (Fig. 1E).

### Relationship among average diffusion coefficient, ligand binding affinity, and G protein activation efficiency of mGluR3

We calculated the *D_Av_* of mGluR3 molecules from the MSD (equations 1 and 2 in Methods). The dose-dependent curves showed the LY341495-induced increase and LY379268-induced decrease of *D_Av_* in the absence of other ligands (Fig. 1F). The LY379268-dependent decrease of *D_Av_* was greater in the presence of 100 nM LY341495, and was significantly suppressed by the addition of 1 μM MNI137 (Fig. 1G). To compare the dose-dependency of *D_Av_* with the ligand-binding affinity of mGluR3, we performed an *in vitro* [^3^H]-LY341495 binding assay (Fig. 1H, I). The EC50 value of the LY341495-induced increase of *D_Av_* (Fig. 1F) was at most half that of [^3^H]-LY341495 binding (Fig. 1H). The IC50 values of the LY379268-induced decrease of *D_Av_* without and with 100 nM LY341495 (Fig. 1F, G) were at most twice that of the competition binding curve between LY379268 and 100 nM [^3^H]-LY341495 (Fig. 1H). These results suggested that the dose-dependency of *D_Av_* corresponded well with the ligand-binding affinity. The effect of MNI137 on mGluR3 could not be measured by the [^3^H]-LY341495 binding assay; no significant difference was observed with and without 1 µM MNI137 (Fig. 1I).

We also measured the G protein activation efficiencies of mGluR3 under the same ligand conditions with an *in vitro* [^35^S]-GTPγS binding assay. mGluR3 showed a high G protein activation even without ligands, and this basal activity was suppressed by LY341495 in a concentration-dependent manner (Fig. 1J). This is consistent with a recent study demonstrating that Cl^−^ binding to the ECD causes high basal activity of mGluR3 (24, 25). Thus, the inverse agonist-induced increase and agonist-induced decrease of *D_Av_* in Fig. 2A reflected the change in the equilibrium between the inactive and active states of mGluR3 molecules on the cell surface. Furthermore, 1 μM MNI137 significantly suppressed the agonist-induced increase of G protein activation efficiency (Fig. 1K), as expected from *D_Av_* of mGluR3 (Fig. 1G).

**Fig. 2.**
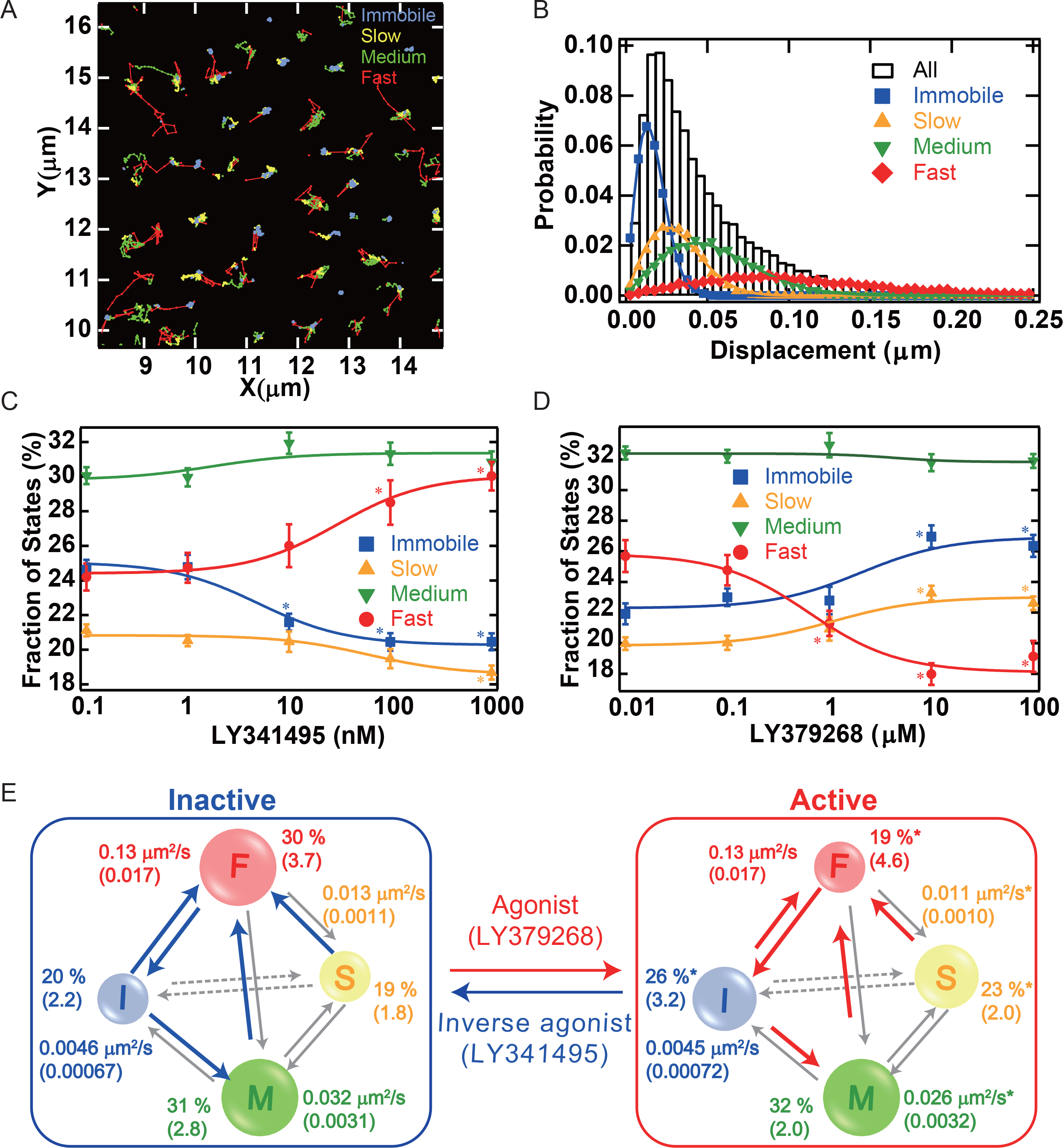
VB-HMM analysis of the trajectories of mGluR3 molecules. **(A)** Every step in the trajectories in Fig. 1B was categorized into four diffusion states. The immobile, slow, medium, and fast states are shown in blue, yellow, green, and red, respectively. **(B)** Histogram of the displacement during 30.5 ms of all the trajectories (open black bars; 28,092 steps from 573 trajectories) on a cell divided into four single-step distributions of random walks (equation 6, Methods). The immobile, slow, medium, and fast states are shown in blue, yellow, green, and red, respectively. **(C, D)** Dose-dependent changes of fractions of the diffusion states. LY341495 dependency is shown in (C). LY379268 dependencies under 100 nM LY314195 are shown in (D). The immobile, slow, medium, and fast states are shown in blue, yellow, green, and red, respectively. All the data are shown as mean ± SEM (*n* = 20 cells). * Significant difference in the fractions compared with the leftmost point in each curve (*p* <0.01; t-test, two-tailed). **(E)** Four state transition diagrams of mGluR3 under inactive (1 μM LY314195) and active (100 μM LY379268 with 100 nM LY341495) ligand conditions. The diffusion coefficient and fraction of each state are shown next to the circles, the size of which reflects the size of the fraction. The SEM is indicated in parentheses (*n* = 20 cells). The arrows between states reflect the rate constants of the state transition estimated from Fig. S4. The significant changes in rate constants between the two conditions in Fig. S4 are shown as colored arrows (inactive: blue, active: red). * Significant difference in the fractions or in the diffusion coefficients compared with the inactive ligand conditions (*p* < 0.01; t-test, two-tailed).

The IC_50_ of LY341495-induced suppression of the basal activity (Fig. 1J) was one order of magnitude smaller than those obtained from the ligand-binding assay (Fig. 1H) and from the dose-dependency of *D_Av_* (Fig. 1F). Furthermore, there was a difference of two orders of magnitude between the EC_50_ values of the LY379268-dependent increase of G protein activation efficiencies with (Fig. 1K) and without 100 nM LY341495 (Fig. 1J), where that with 100 nM LY341495 was similar to those estimated from the ligand binding assay (Fig. 1H) and from the dose-dependency of *D_Av_* (Fig. 1F, G). Generally, it is difficult to estimate the ligand occupancy from a downstream response after amplification of the signaling cascade because the response is usually saturated at a ligand concentration lower than the saturation binding (26). Single-molecule imaging analysis allows us to assess the fraction of receptors in the inactive and active states, which corresponds well to the fraction of ligand binding.

### Ligand-induced changes in the mGluR3 diffusion state distribution

Next, we performed variational Bayesian-hidden Markov model (VB-HMM) clustering analysis (27, 28) to classify the diffusion states of mGluR3. VB-HMM analysis of the total trajectories suggested that the diffusion of mGluR3 molecules could be classified into four states (immobile, slow, medium, fast) (Figs. 2A, B, S3, and Movie 1). The fast and medium states contained transient directional and non-directional movements and their MSD-Δ*t* plots were linear in the average (Fig. S3C, D). In contrast, concave-down MSD-Δ*t* plots were observed in the slow and immobile states (Fig. S3E, F), indicating the confined diffusion of mGluR3 (22). The confinement lengths were estimated to be 140 and 70 nm, respectively (Fig. S3E, F, and Methods), which were consistent with the radii of plasma membrane microdomains (29). The distribution of the apparent oligomer size of mGluR3 in each diffusion state was estimated from the intensity histogram based on the sum of Gaussian functions. The mean intensity of monomeric TMR estimated from the intensity histogram of TMR-labeled CD86, a monomeric membrane protein, on HEK293 cells (11) (Fig. S3G), was about half of that of the highest peak in the histogram of mGluR, suggesting that the majority of mGluR forms dimers (Fig. S3H). The higher-order clusters of mGluR3 were mainly related to the immobile state, where the intensity histogram was right-shifted compared with the other diffusion states (Fig. S3H).

Upon LY341495 stimulation, the fraction of fast state molecules significantly increased, whereas the fractions of immobile and slow state molecules decreased in a dose-dependent manner (Fig. 2C). In contrast, LY379268 stimulation increased the fraction of the immobile and slow states, but decreased the fraction of the fast state (Fig. 2D). To analyze the transitions among the four states, we estimated the time constants of the state transition from the VB-HMM transition array (Figs. 2E, S4). The dose-dependent changes were mainly observed in the transition from the slower to the faster states, suggesting that the activation of mGluR3 made it difficult to escape from the microdomain and that mGluR3 was trapped in a slower state. The diffusion coefficients of medium and slow states estimated from the VB-HMM analysis also changed significantly upon ligand stimulation (Figs. 2E, S5). The ligand-induced changes in *D_Av_* in Fig. 1F, G, were derived from the opposite change in the fraction of the fast state compared with the slow and immobile states, and also from the changes in the diffusion coefficients in the medium and slow states (Fig. 2E).

### Effects of pertussis toxin on mGluR3 molecule diffusion

We analyzed the effect of pertussis toxin (PTX), an inhibitor of G_i/o_ proteins, to link the diffusion state with the G protein-bound state of mGluR3 (Fig. 3A). The PTX treatment decreased the average diffusion coefficient significantly (Fig. 3B-D), reflecting a decrease in the fast state fraction and an increase in the immobile state fraction for the basal (vehicle), inactive (100 nM LY341495) and active (100 μM LY379268) ligand conditions, respectively (Fig. 3H-J). To confirm that the effect of PTX was caused by the loss of the interaction between mGluR3 and G_i/o_, we analyzed the effects of the B oligomer of PTX as a negative control. The B oligomer carries the A protomer that catalyzes ADP-ribosylation of the G_i/o_ α-subunit (30) (Fig. 3A). Treatment with the B oligomer alone did not alter the diffusion of mGluR3 (Fig. 3E-J), indicating that the ADP-ribosylation of the G_i/o_ α-subunit by the A protomer was responsible for the slowing of mGluR3. These results suggest that the fast diffusion state contained G_i/o_ protein-bound mGluR3 for both the inactive and active ligand conditions. This is consistent with previous studies, which demonstrated precoupling of GPCRs with G protein even in the inactive state, enabling fast signal transduction (31–34). The activation of mGluR3 triggered a release of G_i/o_ from the precoupling complex, similar to PTX-induced decoupling (Fig. 3A), thereby decreasing the fast state fraction. Thus, the decrease in *D_Av_* upon agonist stimulation in Fig. 1F, G is partly explained by the decrease in mGluR3 coupling with G_i/o_ protein.

**Fig. 3.**
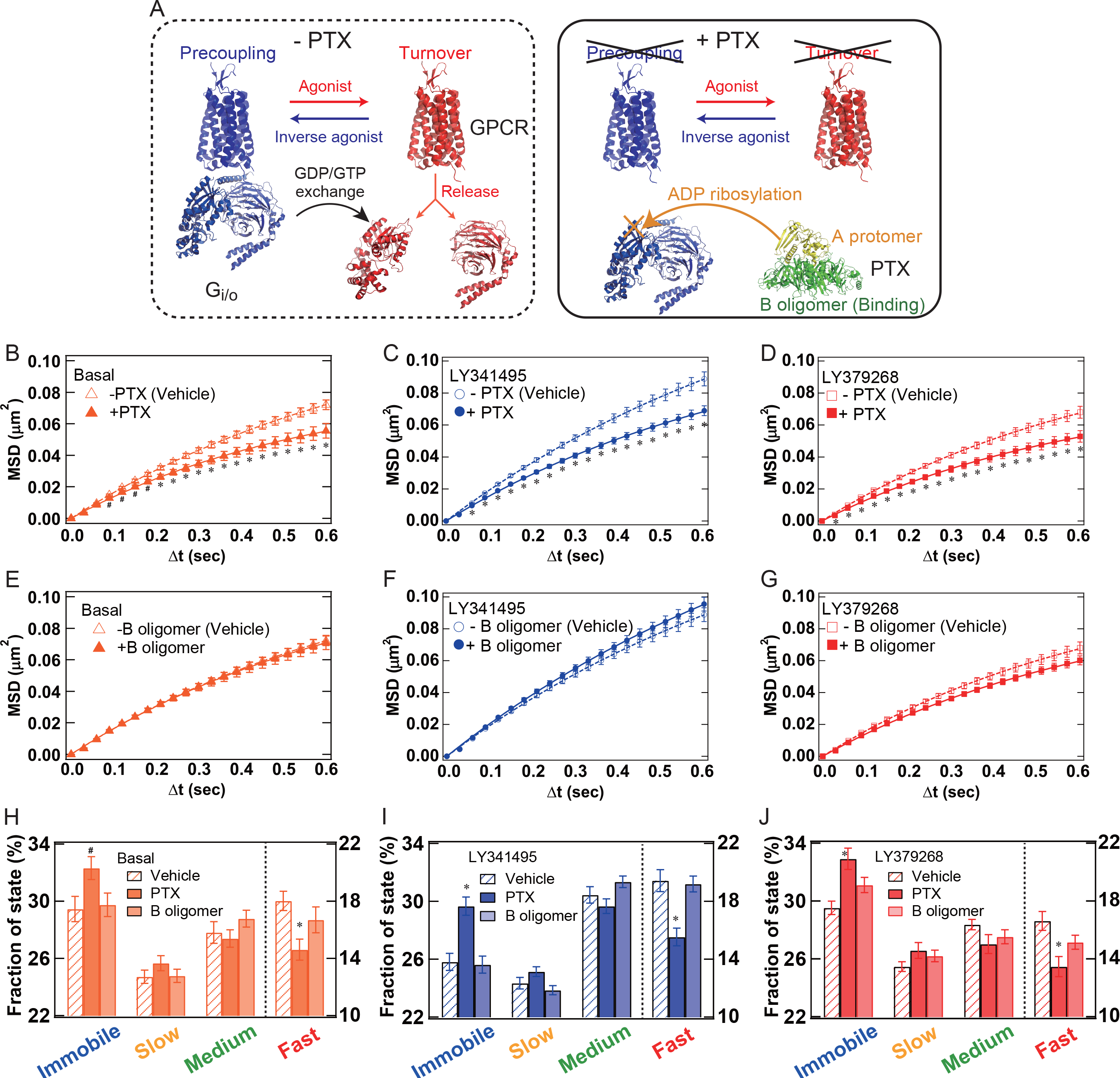
Effects of the PTX treatment on the molecular behavior of mGluR3. **(A)** Schematic model of the effect of PTX on the GPCR/G_i/o_ protein interaction. Without PTX treatment (left panel), a certain amount of GPCR is precoupled with G_i/o_ protein in the inactive state. The G_i/o_ protein is released from GPCR upon activation after the GDP/GTP exchange reaction. GPCR in the active state continuously turns over the G proteins, during which the transient binding and release of G_i/o_ protein occur repeatedly. In contrast, the precoupling and turnover of G_i/o_ protein are inhibited by ADP ribosylation after PTX treatment (right panel). The crystal structures (4OR2, 1GP2, 1GIA, and 1PRT) are drawn with PyMol as representative of GPCR, trimeric G protein, activated Gα, and PTX, respectively. **(B-G)** Comparison of MSD-Δt plots of mGluR3 trajectories with or without PTX. The MSD-Δ*t* plots for the basal (Vehicle), inactive (100 nM LY314195), and active (100 μM LY379268) ligand conditions are shown in (B-D), respectively. Similar comparisons of MSD-Δ*t* plots with or without the PTX B oligomer for the inactive and active ligand conditions are shown in (E-G), respectively. * Significant difference compared with vehicle conditions (*p* < 0.01; t-test, two-tailed). **(H-J)** Comparison of fractions of the diffusion states estimated from VB-HMM analysis. Results from the same experiments in (B-G) are shown as mean ± SEM (*n* = 20 cells). *, ^#^ Significant difference compared with vehicle conditions (*p* < 0.01 and p < 0.05; t-test, two-tailed, respectively).

### Dual-color TIRFM analysis of mGluR3 and G_o_ protein colocalization

To observe the interaction between mGluR3 and G protein directly, we performed dual-color single-molecule imaging analysis. We detected colocalization of TMR-tagged mGluR3 and SiR-tagged G_o_ protein on HEK293 cell membranes in the presence of 1 μM LY341495 (inactive) or 100 μM LY379268 (active) with or without PTX treatment. The rate of SiR dye labeling of G_o_ protein was estimated as ~12% (Fig. S2C).

We observed mGluR3 molecules colocalizing with G_o_ proteins both under the inactive and active ligand conditions (Fig. 4A, B, Movie 2). Binding of G_o_ protein occasionally accelerated mGluR3 diffusion (Movie 2). PTX treatment significantly decreased the MSD of mGluR3 in the presence of exogenous G_o_ protein at saturating ligand concentrations (Fig. 4C, D). The ligand conditions did not affect the probability of mGluR3 and G_o_ protein colocalization; however, PTX treatment significantly decreased it (Fig. 4E). These decreases were caused mainly by the decreased on-rate between mGluR3 and G_o_ proteins, because no significant difference was observed in the cumulative histogram of colocalization duration (Fig. 4F). We compared the diffusion state fraction of mGluR3 colocalized with G_o_ protein (mGluR3/G_o_) with that of mGluR3 colocalized with total mGluR3 (mGluR3/total) in the presence or absence of PTX. Under all conditions, the fractions of the fast and medium states were significantly higher for mGluR3/G_o_ than for mGluR3/total (Fig. 4G, H). Taken together, the PTX-induced deceleration of mGluR3 movement is explained mainly by inhibited formation of the mGluR3/G protein complexes that diffuse faster.

**Fig. 4.**
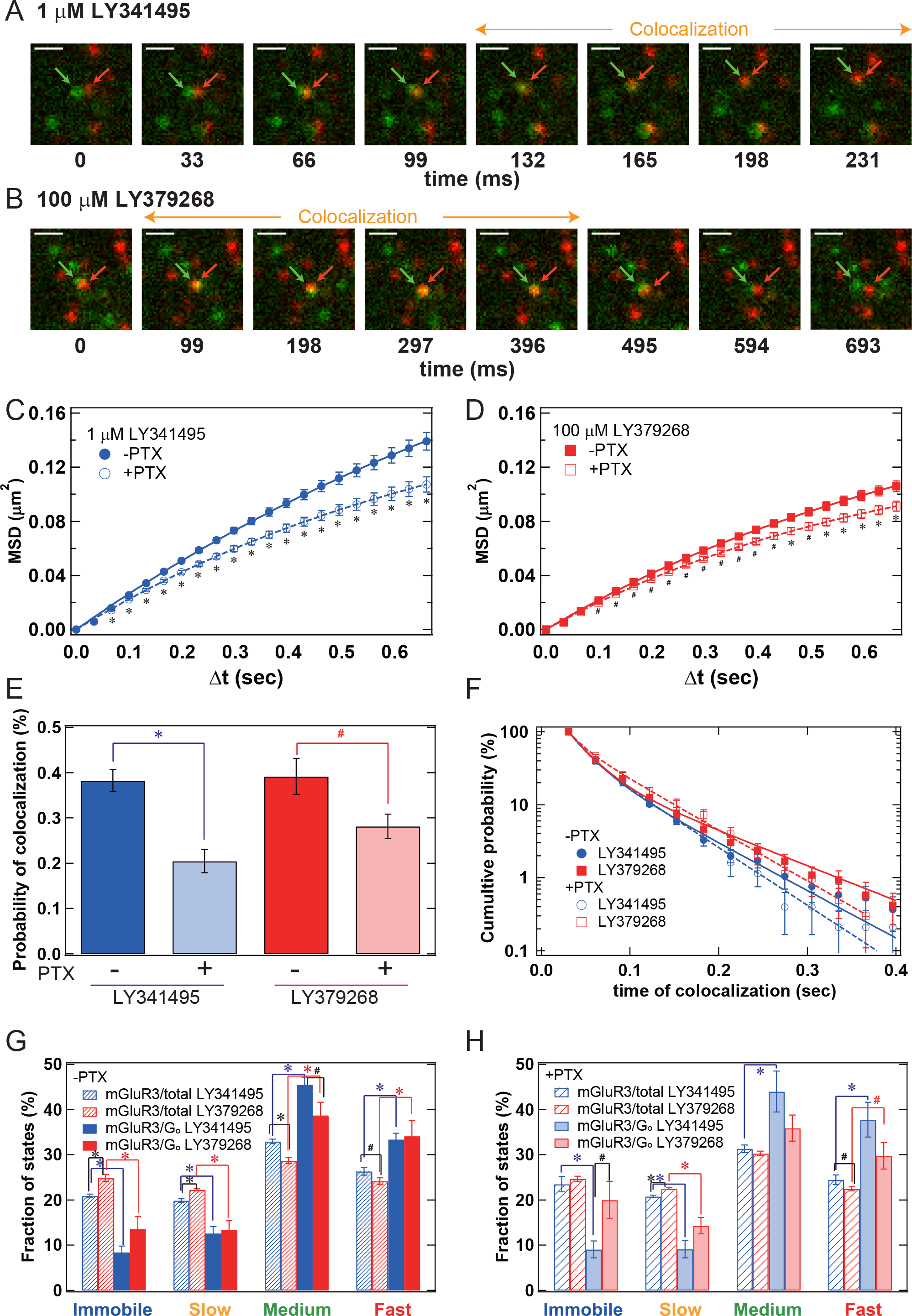
Colocalization analysis of mGluR3 and G_o_ protein. **(A, B)** Representative images of colocalization between TMR-labeled mGluR3 (red) and SiR-labeled G_o_ protein (green). mGluR3 and G_o_ protein formed transient complexes in both the inactive (A: 1 μM LY341495) and active (B: 100 μM LY379268) conditions. Scale bars indicate 1 μm. **(C, D)** Comparison of MSD-Δ*t* plots of the trajectories of mGluR3 in dual-color TIRFM movies with and without PTX. The MSD-Δ*t* plots for the inactive (1 μM LY314195) and active (100 μM LY379268) ligand conditions are shown in (C) and (D), respectively. **(E)** Fraction of mGluR3 colocalized with G_o_ proteins estimated from the total trajectories under the inactive (blue) and active (red) ligand conditions with and without PTX. **(F)** Cumulative probability histograms of the colocalization duration under the inactive (blue) and active (red) ligand conditions without (solid markers) and with (hollow markers) PTX. Curves were fitted using a two-component exponential function markers) (equation 8 in Methods). **(G, H)** Comparison of the fractions of the diffusion states estimated from VB-HMM analysis without (G) and with (H) PTX. The fractions estimated from the trajectories of total mGluR3 molecules under the inactive and active ligand conditions are indicated by blue and red shaded bars, respectively. The fractions estimated from the trajectories of mGluR3 colocalized with G_o_ protein under the inactive and active ligand conditions are indicated by blue and red solid bars, respectively. All data in Fig. 4C-H are shown as means ± SEM (*n* = 20 cells). *^, #^ Significant difference (*p* < 0.01 and p < 0.05; t-test, two-tailed, respectively).

### Dual-color TIRFM analysis of mGluR3 colocalized with clathrin

We investigated the physiological events related to the immobile and slow states that increased upon activation of mGluR3, in contrast to the decrease in the fast state. A TIRFM image showed that the immobile state was related to clustering of mGluR3 molecules followed by internalization (Fig. 5A). Immobile clusters of mGluR3 were formed and disappeared with rapid directional movement (Movie 3). To test whether the clusters were receptors in the CCP, we analyzed the colocalization of TMR-labeled mGluR3 and green fluorescent protein (GFP)-labeled clathrin light chain (CLC) by dual-color TIRFM. When mGluR3 and CLC were colocalized, TMR intensity increased rapidly (Fig. 5B, C, and Movie 4). The intensities of TMR and GFP decreased simultaneously several seconds after colocalization (Fig. 5C). These results suggested that mGluR3 formed a large cluster in a CCP and the cluster was internalized as a clathrin-coated vesicle in the cytoplasmic region, which could not be reached by the evanescent light (Fig. 5A).

**Fig. 5.**
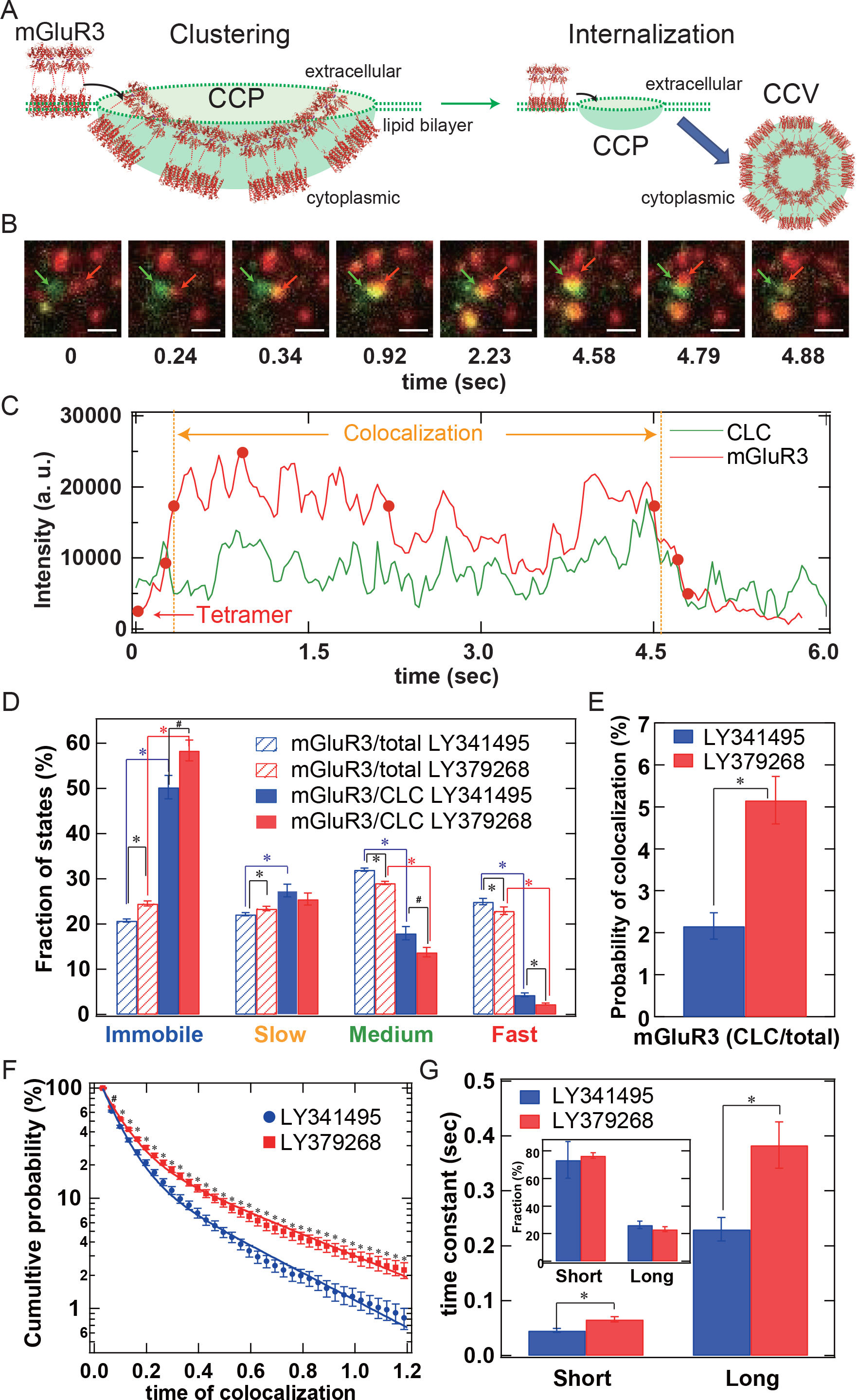
Colocalization analysis of mGluR3 and CLC. **(A)** Schematic of mGluR3 clustering in a CCP followed by internalization. **(B)** Representative images of colocalization of TMR-labeled mGluR3 (red) and GFP-labeled CLC (green). mGluR3 forms a cluster during the colocalization with CLC after 0.34 s and disappears after 4.5 s. Scale bars indicate 1 μm. **(C)** Intensity changes of the particles in B indicated by red and green arrows. A rapid increase in the TMR-labeled mGluR3 intensity coincided with the colocalization with GFP-labeled CLC. TMR and GFP intensities decreased simultaneously after 4.5 s. **(D)** Comparison of the fractions of the diffusion states estimated from VB-HMM analysis. The fractions estimated from the trajectories of total mGluR3 under the inactive (100 nM LY314195) and active (100 μM LY379268) conditions are indicated by blue and red shaded bars, respectively. The fractions estimated from the trajectories of mGluR3 colocalized with CLC under the inactive and active conditions are indicated by blue and red solid bars, respectively. **(E)** Fraction of mGluR3 colocalized with CLC estimated from the trajectories of total mGluR3 under the inactive (blue) and active (red) conditions. **(F, G)** Cumulative probability histograms of the colocalization duration under the inactive (blue) and active (red) ligand conditions. The curves in (F) were fitted using a two-component exponential function (equation 8 in Methods) to show the time constants and fraction (inset) of colocalization in (G). All data in Fig. 5D-G are shown as means ± SEM (*n* = 17 cells for the inactive ligand conditions, 18 cells for the active ligand conditions). *^, #^ Significant difference (*p* < 0.01 and p < 0.05; t-test, two-tailed, respectively).

Next, we quantified the distribution of the diffusion states of mGluR3 colocalized with CLC (mGluR3/CLC) and compared it with the total number of mGluR3 molecules (mGluR3/total) for the inactive (100 nM LY341495) and active (100 μM LY379268) ligand conditions. The immobile and slow state fractions were significantly higher in mGluR3/CLC than in mGluR3/total, indicating that the clathrin binding immobilized the receptor (blue and red lines, Fig. 5D). Comparing the inactive and active ligand conditions demonstrated that the fraction of the immobile state of mGluR3/CLC increased upon activation (black lines, Fig. 5D). Furthermore, the probability and time constant of the colocalization between mGluR3 and CLC were increased significantly after activation (Fig. 5E-G). The cumulative histogram of the colocalization duration was fitted with a double exponential function with short and long time constants (Fig.5F). The time constants were ~1.5 times greater for the active than the inactive ligand condition (Fig. 5G), indicating that the ~1.8-fold increase in the probability of colocalization observed in Fig. 5E was caused mainly by the increased duration of colocalization. Thus, the immobile state fraction in the total trajectories reflected the number of mGluR3 molecules interacting with clathrin molecules, which increased upon activation.

### Effects of RNA interference (RNAi)-mediated CLC inhibition on mGluR3 molecule diffusion

To distinguish between mGluR3 activation and internalization, we measured the effects of RNAi-mediated CLC knockdown on mGluR3 diffusion. Western blotting analysis indicated that transfection of siRNAs specific to CLC reduced the level of CLC by ~50% in HEK293 cells (Fig. 6A). Knockdown of CLC significantly increased the average MSD of mGluR3 under the basal (vehicle) and active (100 μM LY379268) ligand conditions (Fig. 6B, D), reflecting decreases in the slow and immobile states and an increase in the fast state (Fig. 6E, G). In contrast, no significant changes were observed under the inactive (1 μM LY341495) ligand condition (Fig. 6C, F). These results suggested that RNAi of CLC decreased the slowly diffusing mGluR3-clathrin complex and increased the fast state fraction including mGluR3-G protein complex.

**Fig. 6.**
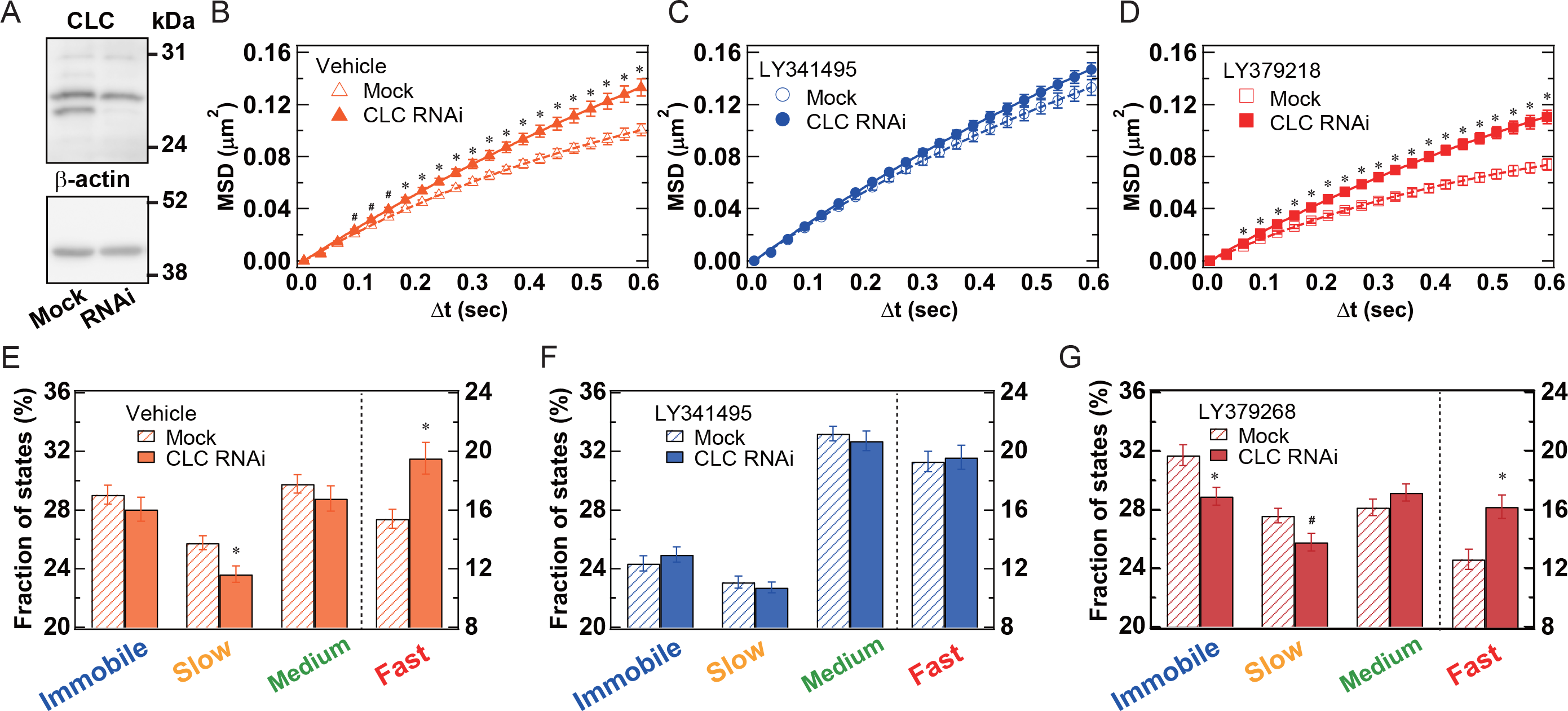
Effects of RNAi of CLC on the molecular behavior of mGluR3. **(A)** Western blot analysis of CLC expression in cells without and with siRNA transfection (Mock and siRNA, respectively). The bands observed 25-30 kDa in the upper panel represent CLC in HEK293 cell lysates. The bands representing β-actin, as a control, in the same lysates are shown in the lower panel. **(B-D)** Comparison of MSD-Δ*t* plots of trajectories of mGluR3 in cells transfected with or without CLC-specific siRNA. The MSD-Δ*t* plots for the basal (Vehicle), inactive (1 μM LY314195), and active (100 μM LY379268) ligand conditions are shown in (B-D), respectively. **(E-G)** Comparisons of the fractions of the diffusion states estimated from VB-HMM analysis. Results from the same experiments in (B-D) are shown as means ± SEM (*n* = 20 cells). *, ^#^ Significant difference compared with vehicle conditions (*p* < 0.01 and p < 0.05; t-test, two-tailed, respectively).

### Correlation between receptor density, mean oligomer size, and D_Av_

We also analyzed the ligand-induced changes in mean oligomer size, which should be related to internalization. However, the mean oligomer size showed no clear dose-dependency (Fig. S6A-C). This may be due to the higher correlation between mean oligomer size and receptor density (Fig. S6D-G). The mean oligomer size of mGluR3 was significantly and positively correlated with receptor density (Fig. S6G). Thus, the strict selection of cells based on receptor density is required to test the ligand effect on oligomer size. In contrast, no significant correlation was observed between receptor density and *D_Av_* (Fig. S7). *D_Av_* is a robust index of mGluR3 activity that is independent of the receptor expression level.

### Generality of the agonist-induced diffusion change of GPCRs

To test the generality of the relationship between the diffusion and activation of GPCRs, we monitored the single-molecule movement of fluorescently labeled GPCRs in other classes (Table 1, Movie 5). Because it is not necessary to label all the molecules in a cell to measure *D_Av_*, we used 30 nM SF650 for labeling to improve the quality of the single-molecule imaging. Under the labeling conditions, ~70% of receptors were labeled with SF650 with lower non-specific binding (Fig. S2B).

**Table 1.** Comparison of *D_Av_* of nine GPCRs in various phylogenetic positions with or without ligands. The class, group, and cluster of GPCRs are listed according to previous reports^1,2^. *D_Av_* was calculated from the MSD in Fig. S8 based on equations 1 and 2 in Methods. All the data are shown as mean ± SEM (*n* = 20 cells). The *p* values of the significant difference between vehicle and ligand conditions were calculated based on Welch’s *t*-test, two-tailed. Under the ligand conditions, GPCR-expressing HEK293 cells were stimulated by the compound listed in the rightmost column.

| GPCR | Class | Group | Cluster | Endogenous ligand | G protein selectivity | $D_{Av}$ ($\mu\text{m}^2/\text{s}$) | | t-test p value | Compounds tested |
| --- | --- | --- | --- | --- | --- | --- | --- | --- | --- |
|  |  |  |  |  |  | Vehicle | Ligand |  |  |
| ADRB2 | A | $\alpha$ | amine | adrenaline | $G_s$ | $0.078 \pm 0.0024$ | $0.051 \pm 0.0023$ | 1.2E-09 | Isoproterenol (10 $\mu\text{M}$) |
| HTR2A | A | $\alpha$ | amine | serotonine | $G_q/G_i$ | $0.079 \pm 0.0022$ | $0.065 \pm 0.0034$ | 2.1E-03 | serotonine (10 $\mu\text{M}$) |
| HRH1 | A | $\alpha$ | amine | histamine | $G_q$ | $0.064 \pm 0.0023$ | $0.045 \pm 0.0025$ | 1.1E-06 | histamine (1 $\mu\text{M}$) |
| ADORA2A | A | $\alpha$ | MECA | adenosine | $G_s$ | $0.066 \pm 0.0022$ | $0.058 \pm 0.0016$ | 8.0E-03 | NECA (10 $\mu\text{M}$) |
| FFAR4 | A | $\alpha$ | melatonin | free fatty acid | $G_q$ | $0.083 \pm 0.0025$ | $0.048 \pm 0.0073$ | 3.5E-13 | DHA (100 nM) |
| CXCR4 | A | $\gamma$ | chemokine | chemokine | $G_i$ | $0.087 \pm 0.0034$ | $0.068 \pm 0.0017$ | 1.7E-05 | CXCL12 (20 nM) |
| F2R | A | $\delta$ | purin | thrombin | $G_q/G_i/G_{12}$ | $0.084 \pm 0.0041$ | $0.060 \pm 0.0017$ | 1.4E-05 | TRAP-6 (10 $\mu\text{M}$) |
| GCGR | B | secretin | peptide | glucagon | $G_s/G_q$ | $0.075 \pm 0.0028$ | $0.042 \pm 0.0025$ | 8.7E-11 | glucagon (1 $\mu\text{M}$) |
| mGluR3 | C | - | amino acid | glutamate | $G_i$ | $0.047 \pm 0.0022$ | $0.039 \pm 0.0018$ | 9.1E-03 | LY379268 (100 $\mu\text{M}$) |
| | | | | | | $0.047 \pm 0.0015$ | $0.064 \pm 0.0018$ | 1.6E-08 | LY341495 (1 $\mu\text{M}$) |

We compared the MSD-Δ*t* plots of the trajectories of GPCR molecules with and without agonist stimulation (Fig. S8), and calculated *D_Av_* as listed in Table 1. All the GPCRs tested showed significant slowing upon agonist stimulation regardless of the phylogenetic positions, chemical properties of the ligands, and G protein-coupling selectivity (Table 1, Fig. S8). In the absence of ligands, mGluR3 showed lower *D_Av_* (0.047 μm^2^/s) than other GPCRs (0.06-0.09 μm^2^/s), which corresponded to that of mGluR3 with 1 μM LY341495 (0.064 μm^2^/s) (Table 1). Thus, the diffusion coefficient of mGluR3 was similar to those of other GPCRs in the inactive state (Fig. 2E). In the presence of an agonist, the *D_Av_* of GPCRs was 0.04-0.07 μm^2^/s. A drug effect on each GPCR was accurately detected by SMT analysis as a change in *D_Av_* (Table 1), but the absolute values of *D_Av_* varied between GPCRs.

## Discussion

The present study provides a new method for assessing the effects of drugs on GPCRs by monitoring the diffusion behavior of GPCRs. We first obtained proof-of-concept of the applicability of single-molecule imaging to the pharmacology of a class C GPCR, mGluR3. The basal activity, agonist-induced activation, inverse agonist-induced inactivation, and NAM-dependent suppression of activity can be evaluated by measuring *D_Av_* of mGluR3 on the living cell surface (Fig. 1). The dose-dependent change of *D_Av_* was derived from the population shift of the four diffusion states and the change of the diffusion coefficient of each state (Fig. 2E). The plasma membrane is partitioned into submicrometer-scale domains with different lipid compositions, including lipid rafts (35). Assuming that the four diffusion states are determined primarily by the lipid environment of receptor molecules, ligand-induced conformational changes followed by interactions with other molecules would alter the accessibility of the receptor to membrane domains. A previous single-molecule imaging study also suggested that the lateral diffusion of transmembrane proteins is transiently anchored by the actin cytoskeleton, which impedes diffusion across the membrane domains freely (36). Furthermore, CCPs on the plasma membrane restrict the receptors to confined areas (37). The present study suggested that the agonist and inverse agonist stimulations increased and decreased, respectively, the distribution of mGluR3 molecules within these microdomains, which was partly related to the binding partner of mGluR3, such as G protein and clathrin (Fig. 7).

**Fig. 7.**
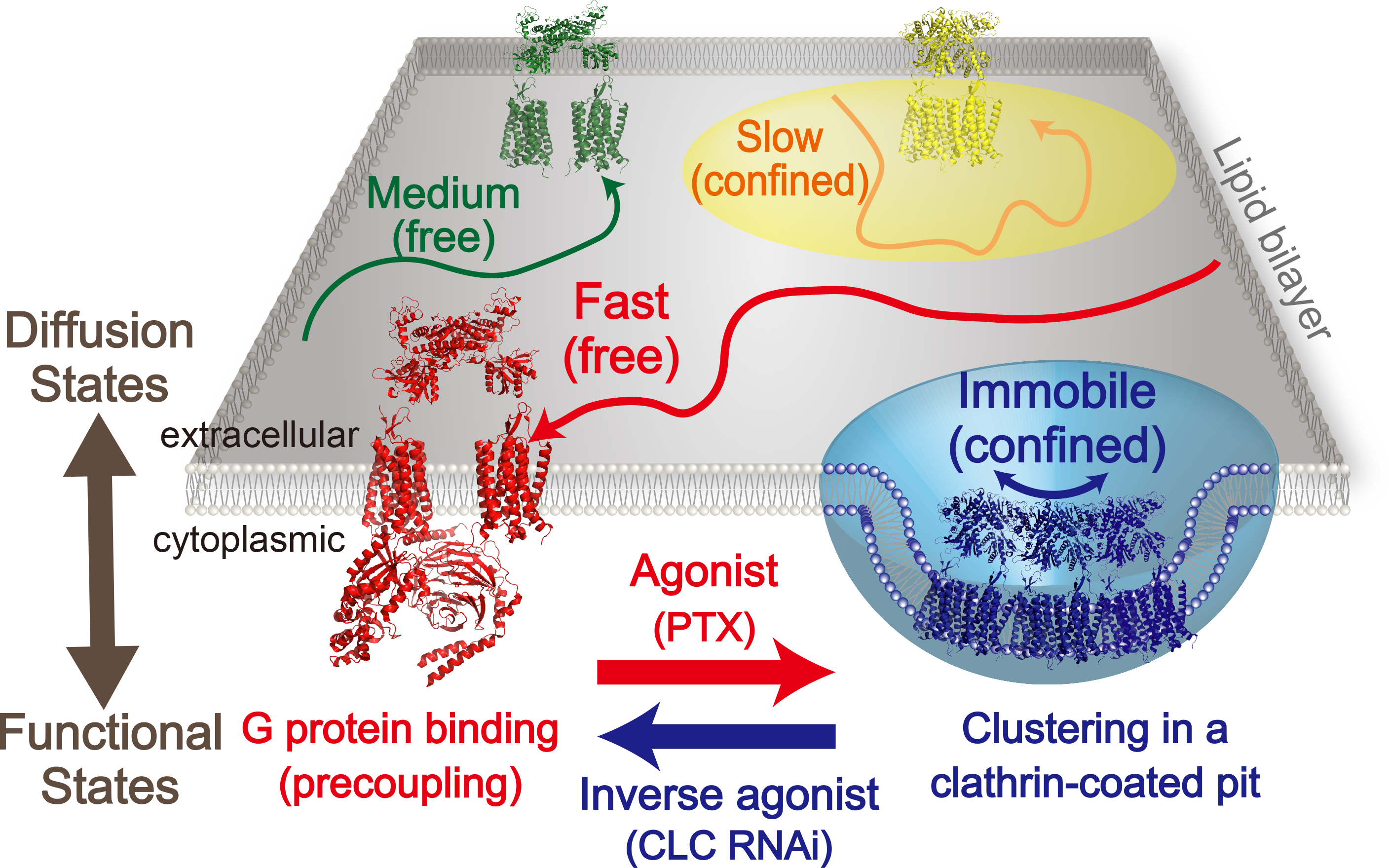
Schematic of mGluR3 diffusion in the plasma membrane. The equilibrium among the four diffusion states of mGluR3 is altered upon ligand stimulation due to differences in the accessibility of mGluR3 to membrane domains that depend on functional states, including G protein binding or clathrin binding domains.

The PTX treatment assay suggested that the fast state is related to the mGluR3 binding with the G protein (Fig. 3, 4). Before the experiment, we expected that PTX would have an effect only under the active ligand conditions; however, this was not the case. The slowing of the mGluR3 by PTX was observed more clearly under the inactive ligand conditions (Fig. 3, 4), providing evidence of mGluR3/G protein precoupling. Furthermore, we directly observed transient formation of the mGluR3/G protein complex under both the inactive and active ligand conditions (Fig. 4A, B, and Movie 2). PTX treatment inhibited formation of the mGluR3/G_o_ complex, which diffuses faster than the average mGluR3 molecule (Fig. 4E, G, H). Although the fraction of fluorescent G_o_-coupled mGluR3 molecules was small (~0.4% of the total mGluR molecules), at least ~3.3% of mGluR3 molecules were colocalized with G_o_, considering the dye labeling rate of G_o_ (~12%) and photobleaching of fluorophores during the measurements. Taking into account the endogenous G_i_ proteins, whose precise concentrations were unknown, the actual fraction of the mGluR3/G_i/o_ protein complex was higher. Thus, the twofold reduction of the mGluR3/G_i/o_ protein complex by PTX treatment seen in Fig. 4E would reasonably explain the ~4% change in the diffusion state distribution of total mGluR3 molecules in Figs. 3 and 4.

Precoupling with G proteins has been demonstrated for class A GPCRs such as adrenergic and muscarinic acetylcholine receptors. The ternary complex model, which assumes that receptor/G protein precoupling induces the high-affinity state of GPCRs, was proposed before the discovery of G proteins (38), and it is widely accepted to depict the properties of agonist/GPCR associations. Although there was no evidence of high-affinity state formation of mGluRs, a previous study using nanodiscs demonstrated that a single TMD of mGluR2, which cannot bind glutamate, can couple with G proteins (39). However, the basal activity of GPCRs, including mGluR3, made it difficult to distinguish the receptor/G protein precoupling complex from spontaneously activated receptor binding to G proteins using biochemical methods. Studies using Förster resonance energy transfer and fluorescence recovery after photobleaching confirmed the presence of precoupling complexes of the _2A_ adrenergic receptor, muscarinic M3 and M4 receptors, adenosine A1 receptor, and protease-activated receptor 1 under the inactive ligand conditions (31–34). In the present study, we could not distinguish the precoupling complex from the active complex by colocalization probability or duration (Fig. 4E, F). From the view point of diffusion, the active complex showed a trend toward becoming immobile compared with the precoupling complex (Fig. 4G, H). This is consistent with a recent dual-color TIRFM analysis of adrenergic receptors and G proteins suggesting that the active receptor/G protein complex is formed in a confined region of the plasma membrane (12). The difference in mobility between precoupling and active complexes may be an additional reason for the increase in the mGluR3 immobile fraction upon activation.

Currently, it is unknown why G protein binding sometimes accelerates the diffusion of mGluR3 (Fig. 4G and Movie2), even though simple physical models predict slower diffusion of particles of larger volume (40). We speculate that the major determinant of the diffusion coefficient of mGluR3 is the viscosity of the membrane surrounding the receptor molecule, which is dependent on the lipid composition. G protein binding would increase the accessibility of mGluR3 to a less viscous membrane environment. In addition, a previous report revealed that crossing the diffusion barrier was controlled by the interaction between the C-terminal region of mGluR5 and a cytosolic partner in astrocytes (41). Receptor/G protein precoupling, in which the C-terminal region of the GPCR also plays an essential role (33), would affect the ability to cross the membrane microdomain for a similar reason.

The recruitment of GPCRs into CCPs is a well-established mechanism for endocytosis regardless of the GPCR family. The dual-color TIRFM analysis demonstrated that the immobile state of mGluR3 is related to the interaction with clathrin molecules (Fig. 5). During desensitization, GPCRs are phosphorylated by G protein-coupled receptor kinases, followed by the recruitment of arrestins (42). Then, the GPCR/arrestin complexes are gathered into CCPs through the interactions between arrestin, clathrin, and the AP2 adaptor (42). Previously, the clathrin-mediated endocytosis of class A GPCRs, including adrenergic and opioid receptors, was analyzed by TIRFM under high-expression conditions where a single receptor molecule could not be resolved, and it was demonstrated that the GPCR cargo regulates the surface residence time of CCPs (43, 44). The present results indicate that the time constant of colocalization between mGluR3 and CLC molecules increases upon agonist stimulation (Fig. 5F, G), which is qualitatively consistent with these previous reports. The absolute values of the colocalization time constant were two orders of magnitude shorter than the previously reported values for bulk imaging, and is probably due to the higher photobleaching rate of single TMR ligands (~3 s) in single-molecule imaging. Thus, it is rare to observe the whole process, from the recruitment of receptors into the CCP to the internalization, as shown in Fig. 5B, C, where the clustering rate of the receptor-clathrin complex was greater than the photobleaching rate. RNAi-mediated knockdown of CLC also suggested that the mGluR3 interacting with clathrin was in the slow and immobile state. These results are consistent with a single-molecule imaging analysis of neurokinin-1 receptor, a class A GPCR (37).

These physiological events, which affect diffusion, are not specific to class C GPCRs. If a drug effect on a GPCR can be estimated from a common change in the diffusion dynamics, we could perform drug assessments of GPCRs without knowing the specific signaling cascade. Therefore, we verified the generality of the diffusion change upon activation in various GPCRs in other classes. Comparison of the diffusion coefficients of GPCRs with and without an agonist demonstrated that the slowing of receptor diffusion upon activation is a general feature of GPCRs irrespective of the signaling pathways downstream of the receptor (Table 1, Fig. S8).

The agonist-induced increases of diffusion coefficients of GABA_B_ (11) and dopamine D_2_ receptors (13) were reported in previous studies. The apparent discrepancy between the present study and previous studies was attributed to the labeling method used in each study as described above. Furthermore, there is a clear difference between the analysis in the present study and that used in the study by Tabor et al., (13) who excluded slow-moving receptors (*D* <☐0.02☐μm^2^/s). If we performed a similar analysis here, the immobile and slow diffusion fractions of mGluR3 would be almost completely filtered out, resulting in a misleading evaluation of *D_Av_*.

In conclusion, the diffusion coefficient is a good index for estimating the drug effects on various GPCRs on a living cell. The present method can be applied to HEK293 cells transiently expressing fluorescently labeled GPCRs because *D_Av_* is hardly affected by variability in the cell surface receptor density (Fig. S7). On the other hand, the mean oligomer size was positively correlated with the receptor density on cell surface (Fig. S6). Further comprehensive dual-color TIRFM analyses are required to assess the broad generality of the state-dependent dimerization or oligomerization of GPCRs in the future.

Because it is possible to quantify the diffusion of any GPCR by using TIRFM, our technique could be useful for drug screening of many GPCRs, including orphan GPCRs, about which little is known. We anticipate that the present study will contribute to the future development of a single-molecule dynamics-based pharmacological method for GPCRs.

## Materials and Methods

### Materials

[^3^H]-LY341495 (1.28 TBq/mmol), LY341495, LY379268, NMI137, and NECA, serotonin were purchased from Tocris Cookson. Isoproterenol, histamine, DHA, CXCL12, TRAP-6, and glucagon were purchased from Santa Cruz, Wako, Sigma Aldrich, Thermo Fisher, BACHEM, and CEDARLANE, respectively. [^35^S]-GTPγS (37 TBq/mmol) was purchased from PerkinElmer Life Sciences. PTX and B oligomer were purchased from Wako Chemicals. Human CD86 cDNA was purchased from OriGene. siRNAs targeting CLCs (5′-GACUUUAACCCCAAGUCUAGC-3′ and 5′ - UAGACUUGGGGUUAAAGUCAC-3′ for CLTA; 5′ - GGCUUAAAGGGUGUGUUGUUG-3′ and 5′-ACAACACACCCUUUAAGCCAA-3′ for CLTB) were designed and purchased from RNAi Inc. The anti-CLC monoclonal antibody (C1985)and anti-β-actin monoclonal antibody (A5441) were purchased from Sigma.

### Construction of cDNA

The HaloTag7 (Promega) coding sequence was amplified by PCR, and was fused at the C-terminus of mouse mGluR3 with an In-Fusion HD Cloning Kit (Clontech). To quantify the expression of wild type and HaloTag-fused mGluR3 by Western blotting analysis, the epitope sequence of the anti-bovine rhodopsin monoclonal antibody Rho1D4 was also fused at the C-terminus. The cDNAs of mGluR3s were introduced into the pcDNA3.1 mammalian expression vector (Invitrogen). The cDNAs of other GPCRs (ADRB2, HTR2A, HRH1, ADORA2A, FFAR4, CXCR4, F2R, GCGR) were purchased from Promega, and the receptor coding sequences were inserted into pFC14K HaloTag CMV Flexi Vector. The CD86 (M1-R277) coding sequence was amplified by PCR, and inserted into the pEGFP-N1 mammalian expression vector (Clontech), where the coding sequence of EGFP was swapped with that of HaloTag7. The SNAP-tag (NEB) coding sequence was amplified by PCR and inserted in the loop region between L91 and G92 of the mouse Gα_o_subunit coding cDNA, as reported previously (12). The SNAP-tagged Gα_o_subunit coding sequence was inserted into the pFC15A HaloTag CMVd1 Flexi Vector without the HaloTag coding sequence to reduce the expression level for single-molecule imaging. The cDNA of GFP-tagged CLC was constructed as previously reported (45).

### Single-molecule imaging

HEK293 cells were cultured in DMEM/F12 containing phenol red (Gibco) supplemented with 15 mM HEPES (pH 7.3), 29 mM NaHCO_3_, and 10% FBS at 37 °C under 5% CO_2_. The plasmid DNA of HaloTag-fused mGluR3 was transfected into HEK293 cells cultured on glass coverslips (Matsunami) on a 60-mm dish 1 day before imaging. Lipofectamine 3000 (Invitrogen) was used for transfection. After 15 min incubation at room temperature, the transfection mixture (plasmid DNA (0.1 μg), P3000 reagent (0.2 μL), Lipofectamine 3000 reagent (2.5 μL), and Opti-MEM (120 μL, Gibco)) was added to cells cultured with DMEM/F12 (3 mL) on a 60-mm dish. For dual-color imaging, the plasmid DNA of GFP-fused CLC (0.02 μg) or SNAP-tag-fused G_o_ protein (0.5 μg) was co-transfected with HaloTag-fused mGluR3. After 3 h incubation at 37 °C under 5% CO_2_, the medium was changed to DMEM/F12 without phenol red (3 mL, Gibco) supplemented with 10% FBS. For the RNAi experiment, siRNAs specific to CLTA and CLTB (36 pmol) were transfected using Lipofectamine RNAiMax (6 μL) the day before transfection of mGluR3 cDNA.

After overnight incubation, the HaloTag-fused mGluR3 was labeled with 300 nM HaloTag TMR ligand (Promega) in DMEM/F12 without phenol red for 15 min at 37 °C under 5% CO_2_. For dual-color labeling of mGluR3 and G_o_ protein, 100 nM SNAP-Cell 647-SiR (NEB) was also added at the same time. For imaging of other GPCRs, we used 30 nM SF650 HaloTag ligand (GORYO Chemical). The HaloTag ligand-treated HEK293 cells on coverslips were washed three times with DMEM/F12 without phenol red (3 mL) in a 60-mm dish. For the inhibitor assay, the cells were treated with 5 nM PTX, 5 nM B oligomer, or vehicle for 6 h at 37 °C under 5% CO_2_ before imaging. Cells were washed twice with PBS and DMEM/F12 without FBS before HaloTag ligand treatment.

The coverslip was mounted on a metal chamber (Invitrogen), and washed five times with Hanks’ balanced salt solution (HBSS; 400 μL, Sigma); with 15 mM HEPES (pH 7.3) and 0.01% BSA, without NaHCO_3_). Ligand (5× concentration) or vehicle solution (100 μL) was added to the chamber with 0.01% BSA/HBSS (400 μL) 10 min before imaging. Single-molecule imaging was performed 10-30 min after ligand (or vehicle) stimulation at room temperature (25 °C). The fluorescently-labeled GPCRs on the basal cell membrane were observed with total internal reflection illumination by using an inverted fluorescence microscope (TE2000, Nikon). The cells were illuminated using a 559 nm, 50 mW laser (WS-0559-050, NTT Electronic) or a 532 nm, 100 mW laser (Compass 315M-100) with an ND50 filter for TMR, with a 488 nm, 30% output power of 200 mW laser (Sapphire 488-200, Coherent) for GFP, or with a 637 nm, 140 mW laser (OBIS 637, Coherent) for SiR and SF650 through the objective (PlanApo 60×, NA 1.49, Nikon) by a dichroic mirror (FF493/574, Semrock) for TMR and GFP, (ET Cy3/Cy5, Chroma) for TMR and SiR, or by a single-band filter set (ET Cy5, Chroma) for SF650. The emission light from TMR and GFP/SiR was split into two light paths by a two-channel imaging system (M202J, Nikon) with a dichroic mirror (59004b, Chroma for TMR and GFP, or FF640-FDi01, Semrock for TMR and SiR) and simultaneously detected by two EM-CCD cameras (ImagEM, Hamamatsu) after passing through band-pass filters (ET525/50m and ET605/70m, Chroma for GFP and TMR, or FF01-585/40 and FF01-676/29 for TMR and SiR). The 4× relay lens was placed before the two-channel imaging system to magnify the image (67 nm/pixel). The fluorescence images were recorded with image software (ImagEM HDR, Hamamatsu or MetaMorph, Molecular Devices) with the following settings: exposure time, 30.5 ms for single-color or 33 ms for dual-color imaging; electron multiplying gain, 200; spot noise reduction, on. The cells were fixed to evaluate the accuracies of the positions of TMR-labeled mGluR3, SiR-labeled G_o_ and GFP-labeled CLC was performed according to a previous method (46). The cells on a coverslip were treated with 4% paraformaldehyde/0.2% glutaraldehyde in PBS for 30 min at room temperature, and they were washed five times with HBSS before imaging.

### Analysis of single-molecule images

The multiple TIFF files (16 bit) were processed by ImageJ as follows.

Background subtraction was performed with a rolling ball radius of 25 pixels, and two-frame averaging of the images was then conducted by the Running_ZProjector plugin (Vale Lab homepage, http://valelab.ucsf.edu/~nstuurman/ijplugins/). The dual-color images were aligned by the GridAligner plugin (Vale Lab homepage) based on an affine transform algorithm. The two channels were calibrated with scattering images of gold particles (60 nm) recorded on the same day. To keep the single-molecule intensity constant across the images, the display range of the brightness and contrast was set as a constant range (minimum: 0, maximum: 1800) followed by image conversion to avi format (8 bit) without compression. SMT analysis was performed with G-count software (G-angstrom) based on a two-dimensional Gaussian fitting algorithm with the following parameters: region of interest size, 6 pixels; Fluorescence limit, 12 arbitrary units; loop, seven times; minimum of 15 frames.

The calculation of the parameters from trajectories, the curve fittings, and the illustrations in the Figures were obtained with Igor Pro 6.36 (WaveMetrix) as follows. The MSD within time *n*Δ*t* of each trajectory was calculated by (22)

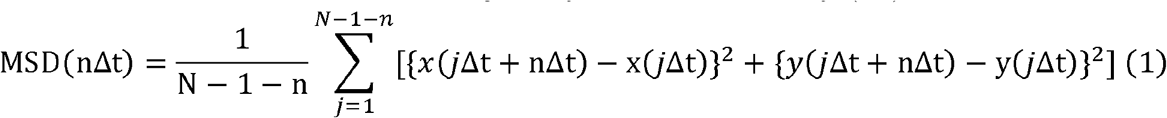

where *n* is the length of frames, Δ*t* is the frame rate (30.5 ms), and *N* is the total frame number of the trajectory. *D_Av_* was calculated based on the two-dimensional diffusion equation

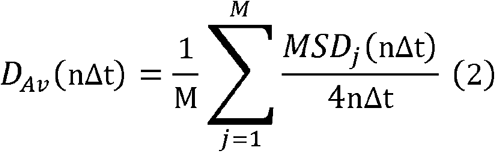

where *MSD*j is the MSD of the *j*-th trajectory and *M* is the total number of trajectories. *D_Av_* in the present study was calculated for *n* = 6 (*n*Δ*t* = 183 ms). The EC_50_ and IC_50_ of the ligand-dependent changes of *D_Av_* were calculated by equations 3 and 4, respectively.

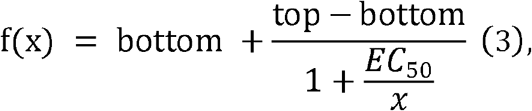

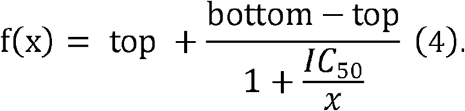

The MSD-Δ*t* plot was fitted by(47)

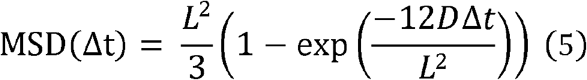

where *L* is the confinement length and *D* is the diffusion coefficient taking the limit of Δ*t* to 0.

The histogram of the displacement (*r* =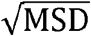) within Δ*t* (30.5 ms) of the trajectories of each HMM diffusion state was fitted by (48)

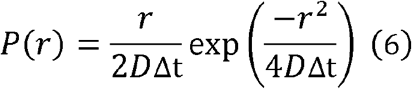

The histogram of the intensity distribution was fitted by sum of the *N* Gaussian

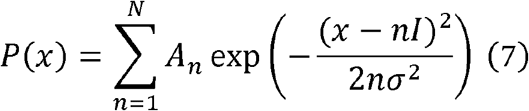

where *n* is the oligomer size, and *I* and σ are mean and SD of a single TMR molecule, respectively. *N* was determined by using the Akaike information criterion. *I* and σ were estimated to be 530 and 210 from the measurement of TMR-labeled CD86. The percentage of each oligomer size, *Percent(n)*, and mean oligomer size on each cell surface were calculated by equations 8 and 9, respectively.

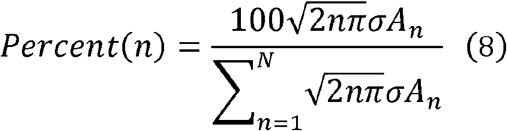

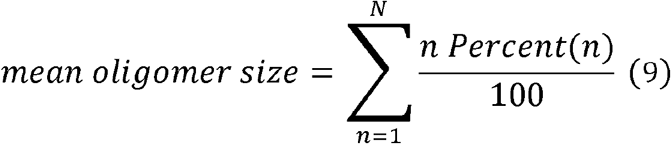

Colocalization between TMR-labeled mGluR3 and SiR-labeled G_o_ protein or GFP-labeled CLC was defined as a distance of less than 100 nm between different proteins in the same frame that are in the same diffusion state, estimated by VB-HMM analysis as described below. It is difficult to distinguish between random colocalization and a true interaction based solely on the distance between two molecules within an image, and the analysis program without the criterion of the proteins being in the same diffusion state defined an interaction as a fast-moving G_o_ protein passing an immobile mGluR3 molecule. It is expected that two molecules are in the same diffusion state if they are truly coupled and moving together. In the VB-HMM analysis of mGluR3, G_o_ protein, and CLC, the single-frame displacement histograms were similarly divided into four diffusion states, whereas the fraction of each state was different depending on the protein (Fig. S9).

The position accuracies of TMR-labeled mGluR3, SiR-labeled G_o_, and GFP-labeled CLC on the fixed cells were estimated to be 28, 24, and 31 nm, respectively, from 1 SD of the displacement distribution of the immobile particles. The error of the alignment between the two channels after the image processing was estimated to be 18 from the difference in the positions of the same gold particles. Therefore, 100 nm corresponded to ~2 SD of the total position accuracy. The time constants of colocalization were estimated from a curve fitting of the cumulative histogram (Figs. 4F and 5F) by the double exponential equation 8.

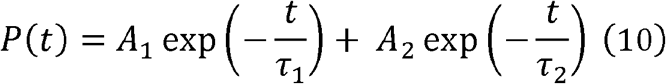

The fraction of two components was estimated from the ratio of A_1_ to A_2_.

### VB-HMM clustering analysis

The VB-HMM analysis was performed with a LabView-based homemade program developed according to previously reported algorithms (27, 28). A trajectory of mGluR3 molecule consists of time series of step displacements. Each time series of the observed data is given as ***X*** = {*x_1_*, …, *x_N_*}, where *N* is the total number of frames. A corresponding time series of diffusion states is defined as ***Z*** = {***z**_1_*, …, ***z**_N_*}. Here, ***z_n_*** ={***z**_n1_*, …, ***z**_nK_*}, in which *K* is the total number of states, and *z_nk_* =1 and 0 otherwise for a molecule in the *k*-th state and *n*-th frame. In order to estimate the state series ***Z*** from the data ***X***, we applied the HMM where ***Z*** is assumed to obey the Markov process with transition matrix ***A***. The distributions of the initial state and transition probability are described as follows, respectively:

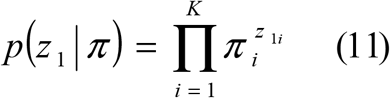

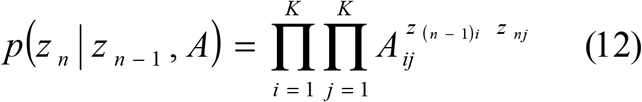

where *π_k_*≡*p(z_li_*=1) satisfies 0 ≤ *i* ≤ 1 and Σ_*i*_π_*i*_ = 1, and *A_ij_* is an element of *A* from the *i* to *j*-th state and satisfies 0 ≤ *A_ij_* ≤ 1. The distribution of the emission probability, which represents the observation probability of the step displacement, is described with parameters *Φ* as follows,

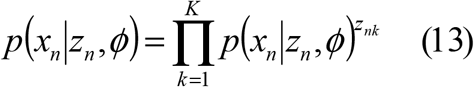

The probability is described with a two-dimensional diffusion equation,

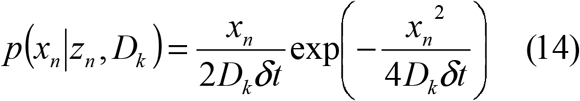

where *D_k_* is the diffusion constant of state *k*, and δ*t* is the frame rate (30.5 ms).

Thus, the joint probability distribution, *p*(***X***, ***Z*** |θ)is

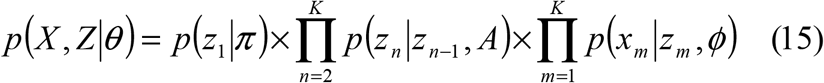

where θ={π, *A, φ*} is the parameters of the observation probability. Molecular states *Z* and model parameters θ were estimated by using the VB method (49) to satisfy the maximum value of the logarithmic likelihood function of *p*(*X*). When the distribution of *θ* is specified by the model *M*,

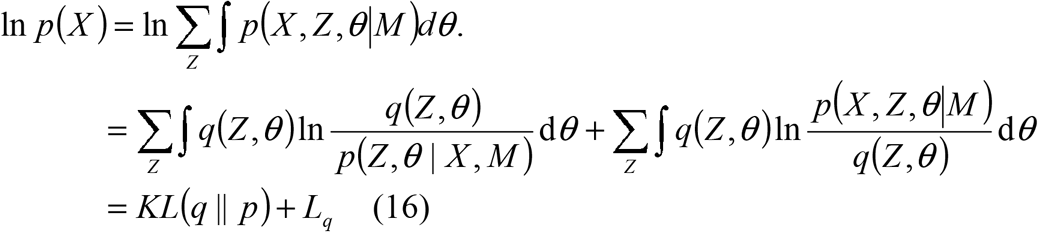

where KL(*q*||*p*) is the Kullback-Leibler divergence between the distribution of model *p* and posterior function *q*. Because KL(*q*||*p*) has fixed values for M and observable *X*, *L_q_* corresponds to the lower bound of ln*P*(*X*). When *q*(***Z***, *θ*) is assumed to be factorized as

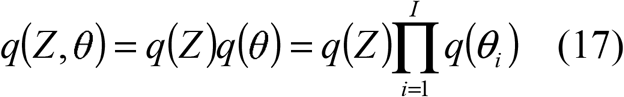

where *I* is the number of parameters. Thus, *L_q_* can be written as,

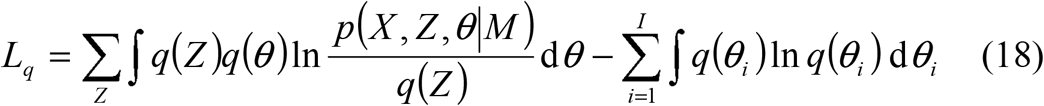

To optimize the distribution functions *q*(*Z*) and *q*(*θ*), the variational Bayesian (VB)-EM algorithm was applied (49). The VB-E step and VB-M step maximize *L_q_* against *q*(*Z*) and *q*(*θ*), respectively. The VB-E step corresponds to the calculation of

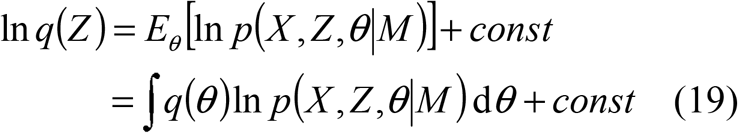

where *E_*θ*_*[…] means the expectation with respect to *θ*. Thus,

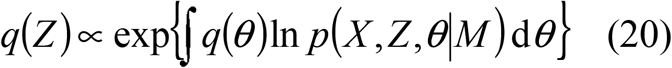

By taking equation (15) into equation (20) and incorporating *M*,

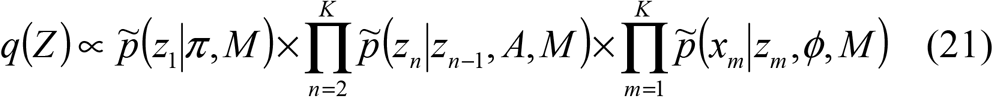

Based on equations (11~13), each term of equation (21) becomes

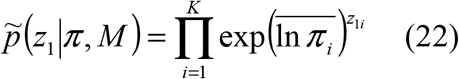

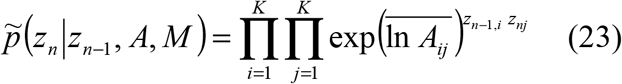

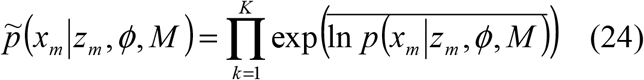

where the overhead lines denote averages. *q*(*Z*) is optimized by the forward-backward algorithm using equations (22~24). Similarly, the VB-M step corresponds to the calculation of

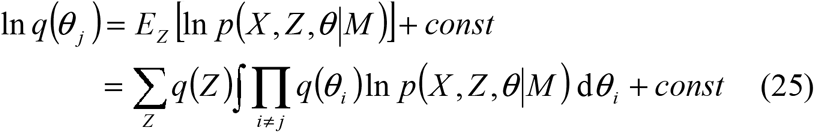

Thus,

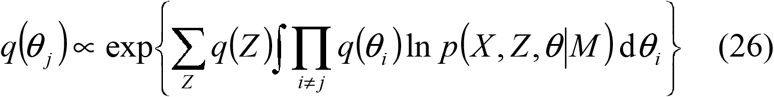

*q*(*θ*_j_) can be factorized to separate terms for each parameter. By optimizing *q*(*θ*_j_), the expectations of parameters are obtained and used as updated values in the next VB-E step.

The Dirichlet distribution was used for given prior functions of the initial state, transition probability distributions, and the calculated posterior functions. Thus, the log of expectation of the initial state is obtained as

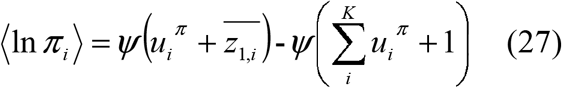

where *Ψ*(*x*) is the digamma function, 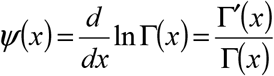 and *u^π^_i_* is the hyper parameter of the prior function and given as a flat probability distribution, *u^π^_i_*=1. The log of expectation of the transition probability is

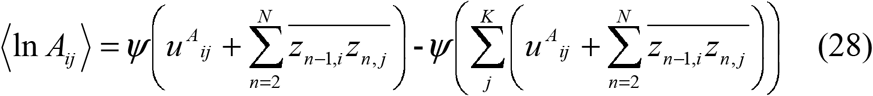

where *u^A^ _ij_* is the hyper parameter and *u^A^ _ij_*=1. For the emission probability of a two-dimensional diffusion equation (equation (14)), the prior function, including the diffusion coefficient (*D_k_*), is given by a gamma distribution, and the log-expectation of parameter *τ*^*D*^_*k*_(=1/2*D_k_δt*) is

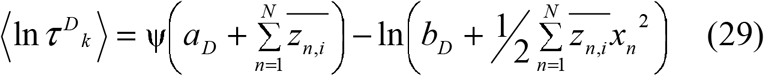

where *a_D_* and *b_D_* are the hyper parameters and assigned values to maximize the lower bound *L_q_*, which was rewritten from equation (18) as

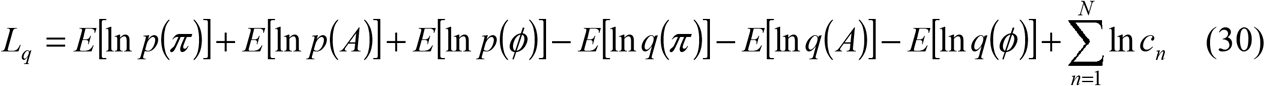

where *c_n_* is the scaling factor calculated in the VB-E step. The iteration between the VBE- and VBM-step is performed until the lower bound converges.

The VB-HMM analysis on the obtained data was carried out using the following procedure: 1) We set the number of states (*N*) and divided the data into *N* groups by the K-means clustering method in which initial values were given by the K means++ method; 2) We calculated the initial parameters of the observation probability for each group; 3) We used the VB-E step to optimize *q*(*Z*) by equations (22~24) with the forward-backward algorithm; 4) We used the VB-M step to update the parameters by equations (27~29); 5) We calculated the lower bound, *L_q_*, by using equation (30) and judging its convergence, except for the first *L_q_*, by determining whether the difference from the previous *L_q_* was less than 0.001%; 6) If *L_q_* was not converged, we repeated the next iteration step by repeating steps 3) to 5); and 7) We optimized the state sequence by choosing a state with the highest probability at every frame. For the calculation in 4), we assigned *u_iπ_*= *u^A^ _ij_*= 1, and different fixed values for *a_D_*, *b_D_* that gave maximum lower bounds for the observed trajectory data.

### Heterologous expression and membrane preparation for in vitro biochemical assay

Heterologous expression of mGluR3 in HEK293 cells for *in vitro* biochemical assay was performed according to previously reported methods (20). The plasmid DNA (10 μg/100-mm dish) of mGluR3 or pCAG vector (mock) was transfected into HEK293 cells growing to ~40% confluency in DMEM/F12 supplemented with 10% FBS by the calcium-phosphate method. The cells were collected 48 h after transfection by centrifugation, and the pellet was washed with PBS (1 mL, pH 7.4). The cell pellet in a 1.5 mL tube was homogenized with a pellet mixer in 50% sucrose in buffer A (50 mM HEPES (pH 6.5) and 140 mM NaCl) prior to centrifugation. The supernatant containing the plasma membrane was diluted in two volumes of buffer A and recentrifuged. The membrane pellet was washed with buffer A and stored at −80 °C.

### Western blotting

The mGluR3 containing membrane pellet was suspended in the sample buffer (62.5 mM Tris-HCl (pH 6.8), 4% SDS, 10% glycerol) with or without 2.5% - mercaptoethanol (ME). After 5.5% SDS-PAGE, the electrophoresed proteins were transferred onto a polyvinylidene difluoride (PVDF) membrane and probed with the Rho1D4 antibody (primary antibody) and the HRP-linked anti-mouse IgG (secondary antibody, Cell Signaling #7076). Immunoreactive proteins were detected using the Amersham ECL prime Western blotting detection reagent (GE) with ImageQuant LAS 500 (GE).

To estimate the efficacy of RNAi of CLC, we transfected the siRNAs using the same protocol as described above for single-molecule imaging. After 2 days, the cells were harvested and homogenized using a pellet mixer. The cell lysate was boiled for 20 min in sample buffer containing 2.5% mercaptoethanol. After subjecting the lysate to 15% SDS-PAGE followed by transfer to a PVDF membrane, CLC and β-actin were probed using anti-CLC and anti-β-actin monoclonal antibodies, respectively. The HRP-linked secondary antibody reaction and detection of immunoreactive proteins were performed as described above.

### [^3^H]-Ligand-binding assay of mGluR3

The cell membranes containing mGluR3 were resuspended in HBSS (with 15 mM HEPES (pH 7.1), without NaHCO_3_ (Sigma)), which are the same buffer conditions as for the single-molecule imaging. [^3^H]-LY341495 binding to membranes was measured at room temperature. The membranes (5 μg total protein) were incubated with 0-1 μM [^3^H]-LY341495 in HBSS for 30 min (final assay volume: 20 μL). After incubation, bound and free radioligands were separated by filtration through a nitrocellulose membrane (0.45 μm HATF, Millipore) using a dot-blotter (FLE396AA, ADVANTEC). The nitrocellulose membrane was washed twice with HBSS (200 μL) and dried for 1 h. The pieces of the nitrocellulose membrane were put in a scintillation cocktail (Ultima Gold, PerkinElmer), and the bound [^3^H]-LY341495 was quantified by a liquid scintillation counter (LS6500, Beckman Coulter). Non-specific binding was measured with the mock-transfected HEK293 cell membrane to show the absence of endogenous mGluR in HEK293 cells. The *K*_d_ was calculated by equation 3, where EC_50_ was replaced by *K*_d_. Displacement by LY379268 of [^3^H]-LY341495 bound to the cell membranes expressing mGluR3 was also measured at room temperature with or without NMI137. A mixture of membranes, 100 nM [^3^H]-LY341495, 0-100 μM LY379268, and 0 or 1 μM MNI137 in HBSS was incubated for 30 min (final assay volume: 20 μL). [^3^H]-LY341495 was quantified as described above. The specific binding was defined using 100 μM LY379268 as the displacer. The IC_50_ values were calculated by equation 4.

### [^35^S]-GTPγS binding assay of mGluR3

The G protein activation efficiencies of mGluR3 were measured under various ligand conditions according to a modified version of our previous method (50). The mGluR3-expressing membrane pellet (final concentration, 11 nM) after sucrose flotation was suspended in 0.02% n-dodecyl-*β*-D-maltopyranoside (DM; Dojindo) in buffer B (50 mM HEPES (pH 6.5), 140 mM NaCl, and 3 mM MgCl_2_) and preincubated with ligands and G_o_ protein (final concentration 200 nM) purified from pig cerebral cortex. After pre-incubation for 30 min at 20 °C, the GDP/GTPγS exchange reaction was started by adding [^35^S]-GTPγS solution. The assay mixture (20 μL) consisted of 50 mM HEPES (pH 6.5), 140 mM NaCl, 5 mM MgCl_2_, 0.01% DM, 0.03% sodium cholate, 5 nM [^35^S]-GTPγS, 500 nM GTPγS (cold), and 500 nM GDP. After incubation for 30 s, the reaction was terminated by adding stop solution (200 μL, 20 mM Tris/Cl (pH 7.4), 100 mM NaCl, 25 mM MgCl_2_, 500 nM GTPγS (cold), and 500 nM GDP) and immediately filtering the sample through a nitrocellulose membrane (0.45 μm HATF, Millipore) to trap [^35^S]-GTPγS bound to G proteins. The nitrocellulose membrane was washed three times with buffer C (200 μL, 20 mM Tris/Cl (pH 7.4), 100 mM NaCl, and 25 mM MgCl_2_) and dried for 1 h. The pieces of the nitrocellulose membrane were put in scintillation cocktail (Ultima Gold, PerkinElmer), and the bound [^35^S]-GTPγS was quantified by a liquid scintillation counter (LS6500, Beckman Coulter). Non-specific binding was measured using the mock-transfected HEK293 cell membrane. The EC_50_ and IC_50_ values were calculated with equations 3 and 4, respectively.

### Saturation binding assay of TMR, SF650, and SiR

The HEK293 cells growing to ~90% confluence on a 100-mm dish were detached using the same protocol for passage, and suspended in DMEM/F12 (4 mL) with 10% FBS. After 15 min incubation at room temperature, the transfection mixture (plasmid DNA of HaloTag-fusion mGluR3, SNAP-tag-fusion G_o_, or pcDNA3.1 vector [2 μg], P3000 reagent [4 μL], Lipofectamine 3000 reagent [5 μL], and Opti-MEM [240 μL, Gibco]) was added to the cell suspension (0.5 mL). After 2 min incubation at room temperature, the lipofectamine-treated cells were diluted with DMEM/F12 (6 mL) with 10% FBS. The cell suspension (100 μL) was added to each well of a black, collagen I-coated 96-well plate (Nunc, Thermo Fisher Scientific). After overnight incubation at 37 °C under 5% CO_2_, the medium in each well was changed to 0-3 μM TMR, SF650, or SiR ligand solution in DMEM/F12 without phenol red (50 μL). After 15 min incubation at 37 °C under 5% CO_2_, the cells were washed three times with DMEM/F12 without phenol red (100 μL), and the medium was finally replaced with 0.001% BSA/HBSS (100 μL) before quantification. Saturation binding of the HaloTag ligand was detected by a microplate reader (FlexStation 3, Molecular Devices) with the following parameters: mode, fluorescence; excitation/cut-off/emission, 530/570/580 nm for TMR, 640/665/675 nm for SF650 and SiR; photomultiplier gain, automatic; flashes per read, 6; read from bottom. The background fluorescence intensity was estimated from the intensity of cells without HaloTag ligand treatment. The non-specific binding was determined by the fluorescence intensity of mock-transfected cells. The specific binding was calculated as the difference between the total binding to cells expressing HaloTag-fusion mGluR3 and the non-specific binding. The data were fitted with the Hill equation

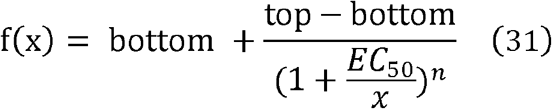

where *n* is the Hill coefficient.

## Supplementary Materials

Fig. S1. Evaluation of the effect of HaloTag fusion to mGluR3

Fig. S2. Comparison of HaloTag and SNAP-tag ligands: affinity, non-specific binding, photostability, fluorescence intensity, and density distribution of mGluR3 molecules on HEK293 cell surfaces in single-molecule imaging

Fig. S3. Example of the VB-HMM analysis of mGluR3 trajectories

Fig. S4. Dose-dependent change in the time constants of the state transition

Fig. S5. Dose-dependent change in the diffusion coefficient of each diffusion state

Fig. S6. Correlation between mean oligomer size and receptor density in various ligand conditions

Fig. S7. Correlation between *D_Av_* and receptor density in various ligand conditions

Fig. S8. MSD-Δt plots of the trajectories of GPCRs with or without agonist

Fig. S9. Histograms showing displacement during 33 ms of the trajectories for mGluR3, G_o_ protein, and CLC molecules, categorized into four diffusion states, using VB-HMM analysis

Movie 1. TIRFM of TMR-labeled mGluR3 molecules on a HEK293 cell

Movie 2. Dual-color TIRFM analysis of TMR-labeled mGluR3 and SiR-labeled G_o_ protein

Movie 3. Accumulation of mGluR3 molecules followed by disappearance with rapid movement

Movie 4. Dual-color TIRFM of TMR-labeled mGluR3 and GFP-labeled CLC

Movie 5. TIRFM of various GPCR molecules on a HEK293 cell with and without agonist

## Acknowledgments

We would like to thank J. Nathans for providing us with the HEK293S cell line, R. S. Molday for the Rho 1D4-producing hybridoma. We would also like to thank T. Yanagida for critical reading of the manuscript. We would like to thank H. Sato for technical assistance.

## Author contributions

M.Y. designed and performed most of the experiments reported. Y. Sako provided guidance throughout. M.H. developed the program of VB-HMM analysis. Y.T. and M.U. upgraded the program of SMT analysis. M.A. designed the siRNAs targeting CLC. M.Y., T.Y., and Y.Shichida constructed cDNAs of mGluR3, and purified G_o_ protein and Rho1D4 antibody. M.M. constructed cDNA of GFP-tagged CLC. M.Y. and Y. Sako wrote the manuscript. M.H., M.U., Y.T., T.Y., and Y. Shichida advised on experiments and manuscript preparation.

## References

1. R. Fredriksson, M. C. Lagerstrom, L. G. Lundin, H. B. Schioth, The G-protein-coupled receptors in the human genome form five main families. Phylogenetic analysis, paralogon groups, and fingerprints. Mol Pharmacol 63, 1256–1272 (2003); published online EpubJun (10.1124/mol.63.6.1256).

2. C. Munk, V. Isberg, S. Mordalski, K. Harpsoe, K. Rataj, A. S. Hauser, P. Kolb, A. J. Bojarski, G. Vriend, D. E. Gloriam, GPCRdb: the G protein-coupled receptor database - an introduction. Br J Pharmacol 173, 2195–2207 (2016); published online EpubJul (10.1111/bph.13509).

3. R. Santos, O. Ursu, A. Gaulton, A. P. Bento, R. S. Donadi, C. G. Bologa, A. Karlsson, B. Al-Lazikani, A. Hersey, T. I. Oprea, J. P. Overington, A comprehensive map of molecular drug targets. Nat Rev Drug Discov 16, 19–34 (2017); published online EpubJan (10.1038/nrd.2016.230).

4. X. L. Tang, Y. Wang, D. L. Li, J. Luo, M. Y. Liu, Orphan G protein-coupled receptors (GPCRs): biological functions and potential drug targets. Acta Pharmacol Sin 33, 363–371 (2012); published online EpubMar (10.1038/aps.2011.210).

5. R. Zhang, X. Xie, Tools for GPCR drug discovery. Acta Pharmacol Sin 33, 372–384 (2012); published online EpubMar (10.1038/aps.2011.173).

6. Y. Sako, S. Minoguchi, T. Yanagida, Single-molecule imaging of EGFR signalling on the surface of living cells. Nat Cell Biol 2, 168–172 (2000); published online EpubMar (10.1038/35004044).

7. Y. Miyanaga, S. Matsuoka, M. Ueda, Single-molecule imaging techniques to visualize chemotactic signaling events on the membrane of living Dictyostelium cells. Methods Mol Biol 571, 417–435 (2009) 10.1007/978-1-60761-198-1_28).

8. M. Hiroshima, Y. Saeki, M. Okada-Hatakeyama, Y. Sako, Dynamically varying interactions between heregulin and ErbB proteins detected by single-molecule analysis in living cells. Proc Natl Acad Sci U S A 109, 13984–13989 (2012); published online EpubAug 28 (10.1073/pnas.1200464109).

9. J. A. Hern, A. H. Baig, G. I. Mashanov, B. Birdsall, J. E. Corrie, S. Lazareno, J. E. Molloy, N. J. Birdsall, Formation and dissociation of M1 muscarinic receptor dimers seen by total internal reflection fluorescence imaging of single molecules. Proc Natl Acad Sci U S A 107, 2693–2698 (2010); published online EpubFeb 9 (10.1073/pnas.0907915107).

10. R. S. Kasai, K. G. Suzuki, E. R. Prossnitz, I. Koyama-Honda, C. Nakada, T. K. Fujiwara, A. Kusumi, Full characterization of GPCR monomer-dimer dynamic equilibrium by single molecule imaging. J Cell Biol 192, 463–480 (2011); published online EpubFeb 7 (10.1083/jcb.201009128).

11. D. Calebiro, F. Rieken, J. Wagner, T. Sungkaworn, U. Zabel, A. Borzi, E. Cocucci, A. Zurn, M. J. Lohse, Single-molecule analysis of fluorescently labeled G-protein-coupled receptors reveals complexes with distinct dynamics and organization. Proc Natl Acad Sci U S A 110, 743–748 (2013); published online EpubJan 8 (10.1073/pnas.1205798110).

12. T. Sungkaworn, M. L. Jobin, K. Burnecki, A. Weron, M. J. Lohse, D. Calebiro, Single-molecule imaging reveals receptor-G protein interactions at cell surface hot spots. Nature 550, 543–547 (2017); published online EpubOct 26 (10.1038/nature24264).

13. A. Tabor, S. Weisenburger, A. Banerjee, N. Purkayastha, J. M. Kaindl, H. Hubner, L. Wei, T. W. Gromer, J. Kornhuber, N. Tschammer, N. J. Birdsall, G. I. Mashanov, V. Sandoghdar, P. Gmeiner, Visualization and ligand-induced modulation of dopamine receptor dimerization at the single molecule level. Sci Rep 6, 33233 (2016); published online EpubSep 12 (10.1038/srep33233).

14. N. Kunishima, Y. Shimada, Y. Tsuji, T. Sato, M. Yamamoto, T. Kumasaka, S. Nakanishi, H. Jingami, K. Morikawa, Structural basis of glutamate recognition by a dimeric metabotropic glutamate receptor. Nature 407, 971–977 (2000); published online EpubOct 26 (

15. R. Vafabakhsh, J. Levitz, E. Y. Isacoff, Conformational dynamics of a class C G-protein-coupled receptor. Nature 524, 497–501 (2015); published online EpubAug 27 (10.1038/nature14679).

16. M. Tateyama, H. Abe, H. Nakata, O. Saito, Y. Kubo, Ligand-induced rearrangement of the dimeric metabotropic glutamate receptor 1alpha. Nat Struct Mol Biol 11, 637–642 (2004); published online EpubJul (

17. M. Yanagawa, T. Yamashita, Y. Shichida, Comparative fluorescence resonance energy transfer analysis of metabotropic glutamate receptors: implications about the dimeric arrangement and rearrangement upon ligand bindings. J Biol Chem 286, 22971–22981 (2011); published online EpubJul 1 (10.1074/jbc.M110.206870).

18. L. Xue, X. Rovira, P. Scholler, H. Zhao, J. Liu, J. P. Pin, P. Rondard, Major ligand-induced rearrangement of the heptahelical domain interface in a GPCR dimer. Nat Chem Biol 11, 134–140 (2015); published online EpubFeb (10.1038/nchembio.1711).

19. C. Brock, N. Oueslati, S. Soler, L. Boudier, P. Rondard, J. P. Pin, Activation of a dimeric metabotropic glutamate receptor by intersubunit rearrangement. J Biol Chem 282, 33000–33008 (2007); published online EpubNov 9 (10.1074/jbc.M702542200).

20. M. Yanagawa, T. Yamashita, Y. Shichida, Glutamate acts as a partial inverse agonist to metabotropic glutamate receptor with a single amino acid mutation in the transmembrane domain. J Biol Chem 288, 9593–9601 (2013); published online EpubApr 5 (10.1074/jbc.M112.437780).

21. B. K. Atwood, J. Lopez, J. Wager-Miller, K. Mackie, A. Straiker, Expression of G protein-coupled receptors and related proteins in HEK293, AtT20, BV2, and N18 cell lines as revealed by microarray analysis. BMC Genomics 12, 14 (2011) 10.1186/1471-2164-12-14).

22. A. Kusumi, Y. Sako, M. Yamamoto, Confined lateral diffusion of membrane receptors as studied by single particle tracking (nanovid microscopy). Effects of calcium-induced differentiation in cultured epithelial cells. Biophys J 65, 2021–2040 (1993); published online EpubNov (10.1016/S0006-3495(93)81253-0).

23. K. Hemstapat, H. Da Costa, Y. Nong, A. E. Brady, Q. Luo, C. M. Niswender, G. D. Tamagnan, P. J. Conn, A novel family of potent negative allosteric modulators of group II metabotropic glutamate receptors. J Pharmacol Exp Ther 322, 254–264 (2007); published online EpubJul (10.1124/jpet.106.117093).

24. A. S. Tora, X. Rovira, I. Dione, H. O. Bertrand, I. Brabet, Y. De Koninck, N. Doyon, J. P. Pin, F. Acher, C. Goudet, Allosteric modulation of metabotropic glutamate receptors by chloride ions. FASEB J 29, 4174–4188 (2015); published online EpubOct (10.1096/fj.14-269746).

25. J. O. DiRaddo, E. J. Miller, C. Bowman-Dalley, B. Wroblewska, M. Javidnia, E. Grajkowska, B. B. Wolfe, D. C. Liotta, J. T. Wroblewski, Chloride is an Agonist of Group II and III Metabotropic Glutamate Receptors. Mol Pharmacol 88, 450–459 (2015); published online EpubSep (10.1124/mol.114.096420).

26. R. R. Neubig, M. Spedding, T. Kenakin, A. Christopoulos, N. International Union of Pharmacology Committee on Receptor, C. Drug, International Union of Pharmacology Committee on Receptor Nomenclature and Drug Classification. XXXVIII. Update on terms and symbols in quantitative pharmacology. Pharmacol Rev 55, 597–606 (2003); published online EpubDec (10.1124/pr.55.4.4).

27. F. Persson, M. Linden, C. Unoson, J. Elf, Extracting intracellular diffusive states and transition rates from single-molecule tracking data. Nat Methods 10, 265–269 (2013); published online EpubMar (10.1038/nmeth.2367).

28. K. Okamoto, Y. Sako, Variational Bayes analysis of a photon-based hidden Markov model for single-molecule FRET trajectories. Biophys J 103, 1315–1324 (2012); published online EpubSep 19 (10.1016/j.bpj.2012.07.047).

29. A. Kusumi, T. A. Tsunoyama, K. M. Hirosawa, R. S. Kasai, T. K. Fujiwara, Tracking single molecules at work in living cells. Nat Chem Biol 10, 524–532 (2014); published online EpubJul (10.1038/nchembio.1558).

30. S. Mangmool, H. Kurose, G(i/o) protein-dependent and -independent actions of Pertussis Toxin (PTX). Toxins (Basel) 3, 884–899 (2011); published online EpubJul (10.3390/toxins3070884).

31. M. Nobles, A. Benians, A. Tinker, Heterotrimeric G proteins precouple with G protein-coupled receptors in living cells. Proc Natl Acad Sci U S A 102, 18706–18711 (2005); published online EpubDec 20 (10.1073/pnas.0504778102).

32. C. Gales, J. J. Van Durm, S. Schaak, S. Pontier, Y. Percherancier, M. Audet, H. Paris, M. Bouvier, Probing the activation-promoted structural rearrangements in preassembled receptor-G protein complexes. Nat Struct Mol Biol 13, 778–786 (2006); published online EpubSep (10.1038/nsmb1134).

33. K. Qin, C. Dong, G. Wu, N. A. Lambert, Inactive-state preassembly of G(q)-coupled receptors and G(q) heterotrimers. Nat Chem Biol 7, 740–747 (2011); published online EpubOct (10.1038/nchembio.642).

34. M. A. Ayoub, E. Trinquet, K. D. Pfleger, J. P. Pin, Differential association modes of the thrombin receptor PAR1 with Galphai1, Galpha12, and beta-arrestin 1. FASEB J 24, 3522–3535 (2010); published online EpubSep (10.1096/fj.10-154997).

35. K. Simons, J. L. Sampaio, Membrane organization and lipid rafts. Cold Spring Harb Perspect Biol 3, a004697 (2011); published online EpubOct 1 (10.1101/cshperspect.a004697).

36. K. Ritchie, R. Iino, T. Fujiwara, K. Murase, A. Kusumi, The fence and picket structure of the plasma membrane of live cells as revealed by single molecule techniques (Review). Mol Membr Biol 20, 13–18 (2003); published online EpubJan-Mar (

37. L. Veya, J. Piguet, H. Vogel, Single Molecule Imaging Deciphers the Relation between Mobility and Signaling of a Prototypical G Protein-coupled Receptor in Living Cells. J Biol Chem 290, 27723–27735 (2015); published online EpubNov 13 (10.1074/jbc.M115.666677).

38. A. De Lean, J. M. Stadel, R. J. Lefkowitz, A ternary complex model explains the agonist-specific binding properties of the adenylate cyclase-coupled beta-adrenergic receptor. J Biol Chem 255, 7108–7117 (1980); published online EpubAug 10 (

39. D. El Moustaine, S. Granier, E. Doumazane, P. Scholler, R. Rahmeh, P. Bron, B. Mouillac, J. L. Baneres, P. Rondard, J. P. Pin, Distinct roles of metabotropic glutamate receptor dimerization in agonist activation and G-protein coupling. Proc Natl Acad Sci U S A 109, 16342–16347 (2012); published online EpubOct 2 (10.1073/pnas.1205838109).

40. M. J. Saxton, Wanted: scalable tracers for diffusion measurements. J Phys Chem B 118, 12805–12817 (2014); published online EpubNov 13 (10.1021/jp5059885).

41. M. Arizono, H. Bannai, K. Nakamura, F. Niwa, M. Enomoto, T. Matsu-Ura, A. Miyamoto, M. W. Sherwood, T. Nakamura, K. Mikoshiba, Receptor-selective diffusion barrier enhances sensitivity of astrocytic processes to metabotropic glutamate receptor stimulation. Sci Signal 5, ra27 (2012); published online EpubApr 3 (10.1126/scisignal.2002498).

42. M. T. Drake, S. K. Shenoy, R. J. Lefkowitz, Trafficking of G protein-coupled receptors. Circ Res 99, 570–582 (2006); published online EpubSep 15 (10.1161/01.RES.0000242563.47507.ce).

43. M. A. Puthenveedu, M. von Zastrow, Cargo regulates clathrin-coated pit dynamics. Cell 127, 113–124 (2006); published online EpubOct 6 (10.1016/j.cell.2006.08.035).

44. A. G. Henry, J. N. Hislop, J. Grove, K. Thorn, M. Marsh, M. von Zastrow, Regulation of endocytic clathrin dynamics by cargo ubiquitination. Dev Cell 23, 519–532 (2012); published online EpubSep 11 (10.1016/j.devcel.2012.08.003).

45. I. Gaidarov, F. Santini, R. A. Warren, J. H. Keen, Spatial control of coated-pit dynamics in living cells. Nat Cell Biol 1, 1–7 (1999); published online EpubMay (10.1038/8971).

46. K. A. K. Tanaka, K. G. N. Suzuki, Y. M. Shirai, S. T. Shibutani, M. S. H. Miyahara, H. Tsuboi, M. Yahara, A. Yoshimura, S. Mayor, T. K. Fujiwara, A. Kusumi, Membrane molecules mobile even after chemical fixation. Nature Methods 7, 865–866 (2010); published online EpubNov (10.1038/nmeth.f.314).

47. Z. Xiao, X. Ma, Y. Jiang, Z. Zhao, B. Lai, J. Liao, J. Yue, X. Fang, Single-molecule study of lateral mobility of epidermal growth factor receptor 2/HER2 on activation. J Phys Chem B 112, 4140–4145 (2008); published online EpubApr 3 (10.1021/jp710302j).

48. K. M. Wilson, I. E. Morrison, P. R. Smith, N. Fernandez, R. J. Cherry, Single particle tracking of cell-surface HLA-DR molecules using R-phycoerythrin labeled monoclonal antibodies and fluorescence digital imaging. J Cell Sci 109 (Pt 8), 2101–2109 (1996); published online EpubAug (

49. M. Beal, (2003).

50. M. Yanagawa, T. Yamashita, Y. Shichida, Activation Switch in the Transmembrane Domain of Metabotropic Glutamate Receptor. Molecular Pharmacology 76, 201–207 (2009); published online EpubJul (10.1124/mol.109.056549).

